# Image quality metrics fail to accurately represent biological information in fluorescence microscopy

**DOI:** 10.1101/2025.08.05.668508

**Authors:** Ihuan Gunawan, Richard J Marsh, Nandini Aggarwal, Erik Meijering, Susan Cox, John G Lock, Siân Culley

## Abstract

Image processing methods offer the potential to improve the quality of fluorescence microscopy data, allowing for image acquisition at lower, less phototoxic illumination doses. The training and evaluation of such methods is informed and driven by full-reference image quality metrics (IQMs); however, these metrics derive from applications to natural scene images, not fluorescence microscopy images. Here we investigate the response of IQMs to common properties of fluorescence microscopy data and whether IQMs are capable of reporting the biological information content of images. We find that IQM scores are biased by image content for both raw and processed microscopy data, and that improvements in IQM values reported after processing are not reliably correlated with performance in downstream analysis tasks. As common IQMs are unreliable proxies for guiding image processing developments in biological fluorescence microscopy, image processing performance should be benchmarked according to downstream analysis success.

---

Fluorescence microscopy is a key quantitative method in biology, allowing for investigation of cellular phenomena with high spatiotemporal resolution. Extracting accurate quantitative information from fluorescence microscopy data requires high quality images (1). However this typically requires imaging at high illumination doses which can be damaging to samples (2). As a result, there has been substantial development in the field of image restoration for images acquired at low illumination doses which intend to improve image quality post-acquisition (3). Similarly, diverse image augmentation approaches, including virtual labelling (4) and computational super resolution (5), aim to further extract biological information in microscopy. Regardless of the image processing method, a key question must be answered: are the generated images faithful to the underlying biology present in the data?

Recent developments in image processing for fluorescence microscopy have been near-universally benchmarked using full-reference image quality metrics (IQMs). The most commonly used metrics are peak signal-to-noise ratio (PSNR (6)), structural similarity index measure (SSIM (7)) and Pearson’s correlation coefficient (PCC (8)) (Supplementary Note 1). PSNR was originally developed for use in signal processing before its extension to images, and SSIM was developed to reproduce key components of the human visual system when perceptually assessing the quality of natural scene images. PCC is not specific to images, and is a general statistical measure used for quantifying the linear correlation between two variables. Importantly, none of these IQMs were developed specifically for fluorescence microscopy data (and indeed, the original SSIM paper explicitly highlights a lack of testing on biomedical images (7)). Fluorescence microscopy images have distinct statistical properties and downstream applications, and the use of IQMs when applied to such data remains chronically under-examined and unvalidated (9, 10). Furthermore, concerns have grown regarding the appropriate use of these metrics across multiple contexts (3, 10).

As image quality has become synonymous with biological resolution, IQMs are currently used as a proxy for comparing the information content or suitability for downstream analysis between ‘test’ images (e.g, raw low illumination dose images or the outputs of image processing) by typically defining quality by their similarity to matched ‘reference’ images (e.g. high illumination dose images) which are presumed to optimally represent underlying biological information (9, 10) (Supplementary Figure S1). This presumed correspondence between image quality estimates and downstream biological analysis accuracy remains without evidential foundation. This is a striking gap in the field, given that IQMs are an underpinning technology guiding both optimisation and evaluation of image processing methods.

Here, we investigate the performance of IQMs in accurately evaluating the quality and biological information content of raw and processed fluorescence microscopy images. We first ask whether IQMs reliably define image quality when assessing raw vs. processed fluorescence images. Second, we ask whether IQM values reliably predict the performance of downstream analysis that extract biological information from images. Together, these approaches give insight into whether current IQMs are useful for assessing the quality of fluores-cence images, and whether they are suitable tools to guide optimisation of image processing methods.

## Results

### Image quality metrics are unreliable for assessing both raw and processed microscopy data

Here, we used PSNR, SSIM and PCC as exemplar IQMs for testing applicability to fluorescence microscopy data. These metrics are commonly used and represent three distinct approaches to assessing image quality. While other metrics are seen in the literature, these are usually closely related to one of these; for example, mean-squared error is a component of PSNR calculation, and variations of SSIM all have the original SSIM metric at their core (Supplementary Note 1). A straightforward evaluation indicates that these IQMs do indeed delineate image quality in fluorescence microscopy data. For a single field of view imaged at several illumination doses, all IQM scores correctly ranged levels of signal (Figure 1a, Supplementary Figure S2a). However, fields with distinct fluorescence coverage (modulated by cell density, size, and marker identity) were scored differently, biasing all IQMs according to sparsity (Figure 1b, Supplementary Figure S2b) and marker identity (Figure 1c). Additionally, IQM score could also fail to reflect both subjective visual quality and objective quality deterministically defined by illumination dose (Figure 1d, Supplementary Figure S2c). Absolute IQM values therefore cannot be interpreted as indicative of quality, given that changes in image structure and sparsity allow higher scores to be achieved by poorer quality images.

**Fig. 1.**
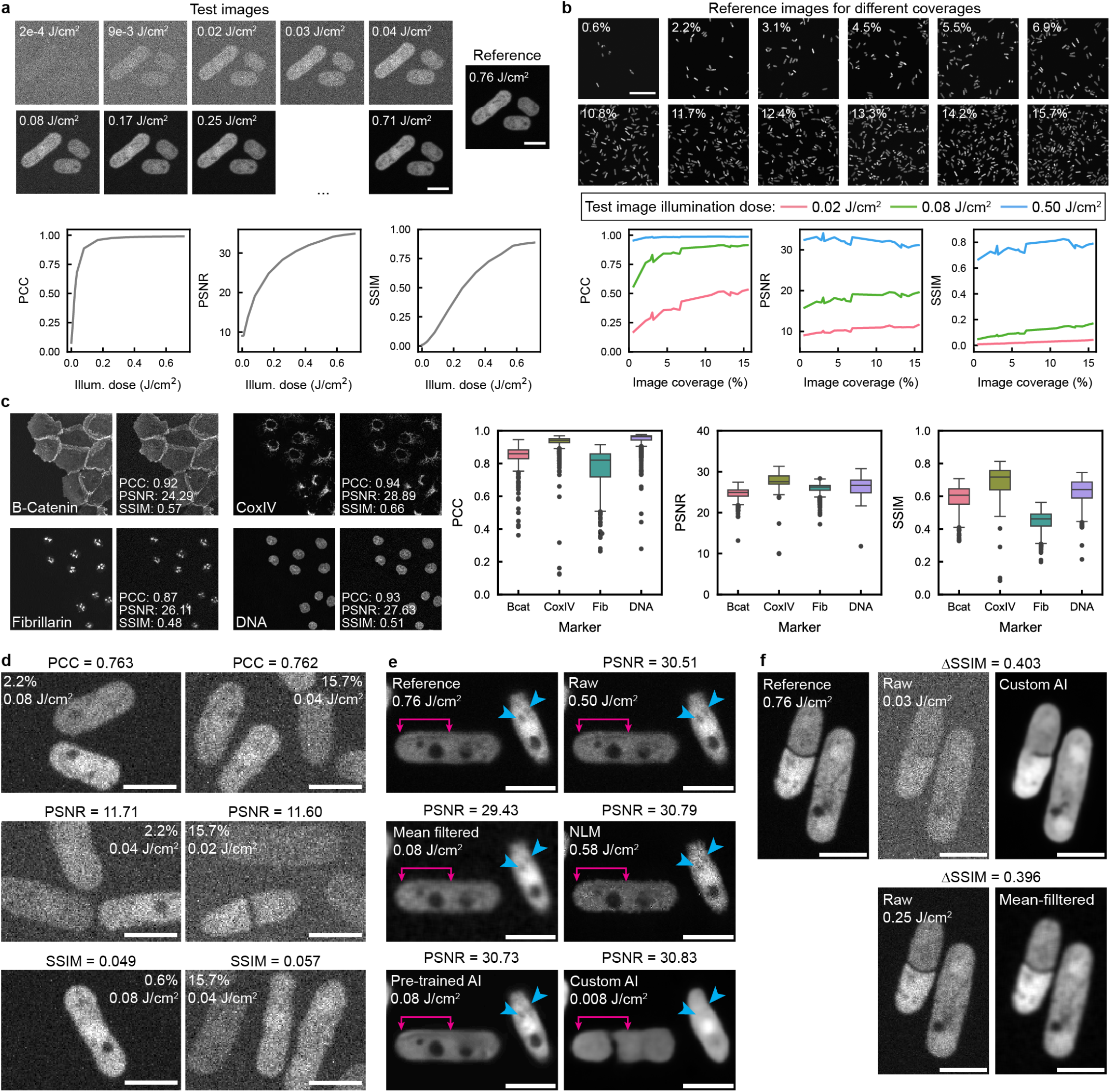
Quality metric behaviour on raw and processed microscopy data. **a** Top: Crops from a series of images of *S. pombe* expressing cytoplasmic sfGFP acquired at a range of illumination doses. Bottom: IQM values for one whole field of view as a function of illumination dose. **b** Whole fields of view of *S. pombe* containing different image coverages, acquired at 0.76 J/cm^2^. Bottom: IQM values plotted as a function of image coverage, for three different test illumination doses. **c** Images of four different subcellular labels in DU145 cells (left images) and the results of adding synthetic noise (right images, with IQM values shown for each image). Box and whisker plots show IQM values for 583 different images of each structure, each after applying the same amount of synthetic noise. Significance tests shown in Table S1. **d** Crops from images of *S. pombe* where the whole fields of view have the same IQM value (top: PCC, middle: PSNR, bottom: SSIM) despite acquisition with difference illumination doses. **e** Crops from images of *S. pombe*, either raw or following image processing, where each whole field of view has PSNR ∼ 30. Annotations indicate fine details whose visibility varies substantially between crops. **f** Crops from images before and after image processing, where for both images processing has improved SSIM by ∼ 0.4. In the top panel, this corresponds to a substantial improvement in visual quality, whereas in the bottom panel similar features are already visible both before and after processing. Scale bars: a - 5 µm, b - 50 µm, d - 5 µm, e - 5 µm, f - 5 µm.

This is particularly relevant when using IQMs to evaluate virtual labelling performance for difference biomarkers, when evaluation of absolute IQM values may be taken to suggest that some labels have been better predicted than others (11). Yet, different markers have fundamentally different scoring ranges given the same underlying quality degradation (Figure 1c), confounding this interpretation, as previously implied for image analysis validation metrics (12). These failings were replicated in bulk analyses (using aggregated scores), which is standard practice in benchmarking (Supplementary Figure S3). Overall, in the presence of Poisson/Gaussian noise, IQMs reliably ranked the levels of signal within a dataset when the same type of structure was imaged with similar coverage in different images, but if either of these factors are not completely controlled, they can become dominant and render the absolute value of the IQM meaningless. This can seriously limit reliable comparison of scores, particularly between studies with distinct underlying data.

To understand the performance of IQMs in assessing image processing techniques, we characterised IQM behaviour across image restoration methods of varying complexity: mean-filtering, non-local means filtering (13) (NLM), and AI-based image restoration using both pre-trained (14) (Denoise.AI) and custom-trained (15) (content aware image restoration, CARE) networks. We applied each method to several datasets, which included variation in acquisition or synthetically modulated signal. Visually distinct images could yield very similar IQM scores (Figure 1e, Supplemen-tary Figure S2d). Likewise, raw and denoised image pairs achieving similar IQM score improvements displayed a variety of different structures (Figure 1f, Supplementary Figure S2e), suggesting that IQMs provide ambiguous and/or inconsistent definitions of visual quality in the present of complex denoising transformations.

### Image quality metrics inconsistently reflect biological information content

Unlike our strictly noise-dependent experiments (across Figure 1a-d) where imaging quality was deterministically controlled by illumination dose, the true quality of computationally processed images is hard to define. One of the most important tasks involving fluorescence microscopy data is the downstream extraction of quantitative biological information via image analysis. It is therefore reasonable to assume that image restoration approaches should aim to enhance the performance of these analyses.

To test whether IQM values of processed images are a dependable proxy for downstream biological data analysis, we considered three sample image analysis tasks: nucleus segmentation, local filament alignment, and whole-image protein localisation classification. In all cases, task performance correlated well with increasing raw data quality as dictated by acquisition parameters (Figure 2a). Thus, higher quality images do indeed enable downstream quantitative analysis results which are closer to the true underlying biology. To systematically assess whether IQM scores for denoised images are indicative of both image quality and of the biological information accessible in a dataset, we compared image processing-induced changes in IQM scores to resulting changes in the accuracy of downstream analyses (Figure 2b). Changes in IQM scores from denoising did not reliably correspond to changes in quantifiable downstream analysis performance, consistent with subjective examples (Figure 1e-f). For segmentation (Figure 2c, Supplementary Figure S4), denoising had no clear impact on performance despite improving IQM scores significantly. This undercuts the assumption that higher IQM scores indicate higher image quality and thus a better or more biologically informative image. Assessments of filament alignment and localisation classification did suggest some correlation between denoising-induced IQM changes and changes in analysis accuracy (Figure 2d- e, Supplementary Figures S5, S6). Yet even in these contexts, IQM scores remain highly ambiguous and thus unreliable, since a given IQM score improvement could correspond with widely dispersed change in analysis performance. For example, note the contradictory performances of segmentation and filament alignment analyses for denoised images with ΔPSNR∼10. Additionally, for segmentation and protein localisation classification in particular, denoising delivered negligible improvements in biological analysis performance compared to better raw data yielding similar IQM score changes (Supplementary Figure S7). In these cases, even a very modest increase in raw data quality generally led to a much greater improvement in analysis accuracy than applying image processing could produce. Thus, any relationships between variable IQM scores and variable biological analysis performance do not generalise between raw and processed images – a critical shortfall given the widespread use of IQMs as benchmarks for computational microscopy (16). In this context, the small and inconsistent effects on downstream performance of sizeable IQM score changes imply that the often incremental IQM score changes reported in many benchmarking studies are unlikely to reflect trustworthy and meaningful changes in the downstream performance of biological analysis tasks.

**Fig. 2.**
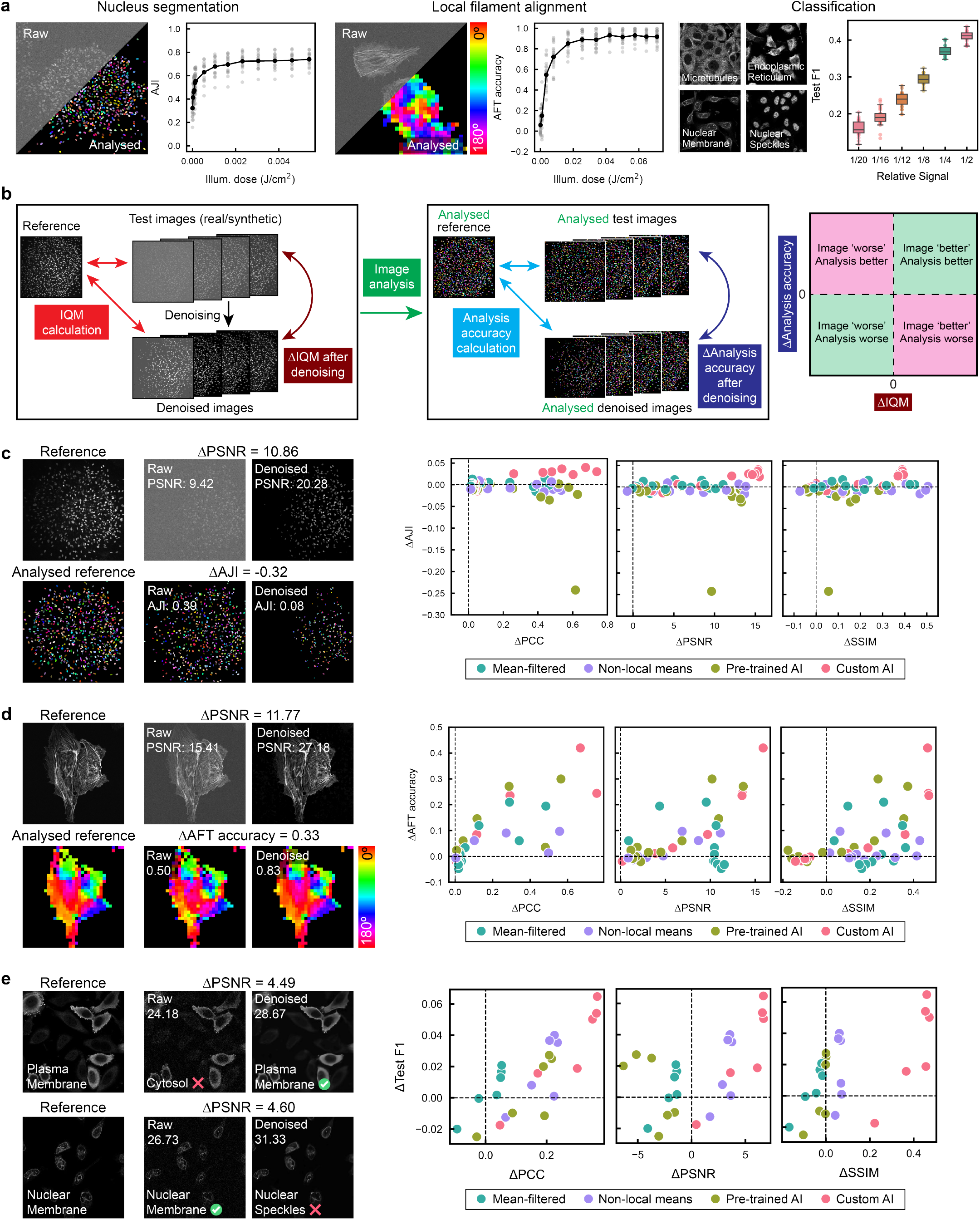
Exploring the relationship between quality metrics and image analysis accuracy. **a** The accuracy of three different analysis tasks - 2D nucleus segmentation in spheroids, AFT filament alignment analysis of actin, and classification of image content - depends on the illumination dose (for real data) and relative signal (for synthetic data). **b** Workflow for exploring whether the changes in IQM values caused by image processing result in improved downstream analysis accuracy. **c** Example reference image and analysis, raw image and analysis, and denoised image (pre-trained AI) and analysis for spheroid segmentation. ΔAJI is plotted as a function of ΔIQM for image grouped by illumination dose for each denoising method. Each point is the mean of 67 different fields of view across four experimental timepoints. **d** Example reference image and analysis, raw image and analysis, and denoised image (pre-trained AI) and analysis for AFT. Plots are displayed as for c; each point is the mean of 20 different fields of view. **e** Example reference images and ground truth classification, and two examples of raw/denoised images with their classification results. Plots are as for c and d, except points here are grouped by the six different synthetic noise levels generated. Each point is the mean test score from five cross-validated folds, each containing 640 test images.

### Image quality metrics can yield totally incorrect results

The results in Figures 1 and 2 highlight important shortcomings and biases of commonly used IQMs in typical benchmarking applications. Yet beyond these shortcomings, we also found evidence that IQMs could produce entirely misleading results. In Figure 3 and Supplementary Figure S8, we show that IQMs can score low-noise images of a completely different cellular structure more highly (i.e. as more similar) than high-noise images of an identically matched cellular structure. For the example images in Figure 3a, when using a high-quality nucleus image as the references, test images of both actin and mitochondria substantially out-score a lower-quality image of the reference nucleus for both PSNR and SSIM. This result is repeatable across multiple fieldsof-view, and for poorer-quality nucleus test images PCC can also be out-scored by an incorrect biological structure (Figure 3b). These results are particularly pertinent in cell biology applications, since most microscopy studies value the accurate representation of spatially distributed molecular labelling as highly as, if not more highly than, an accurate representation of signal intensity.

**Fig. 3.**
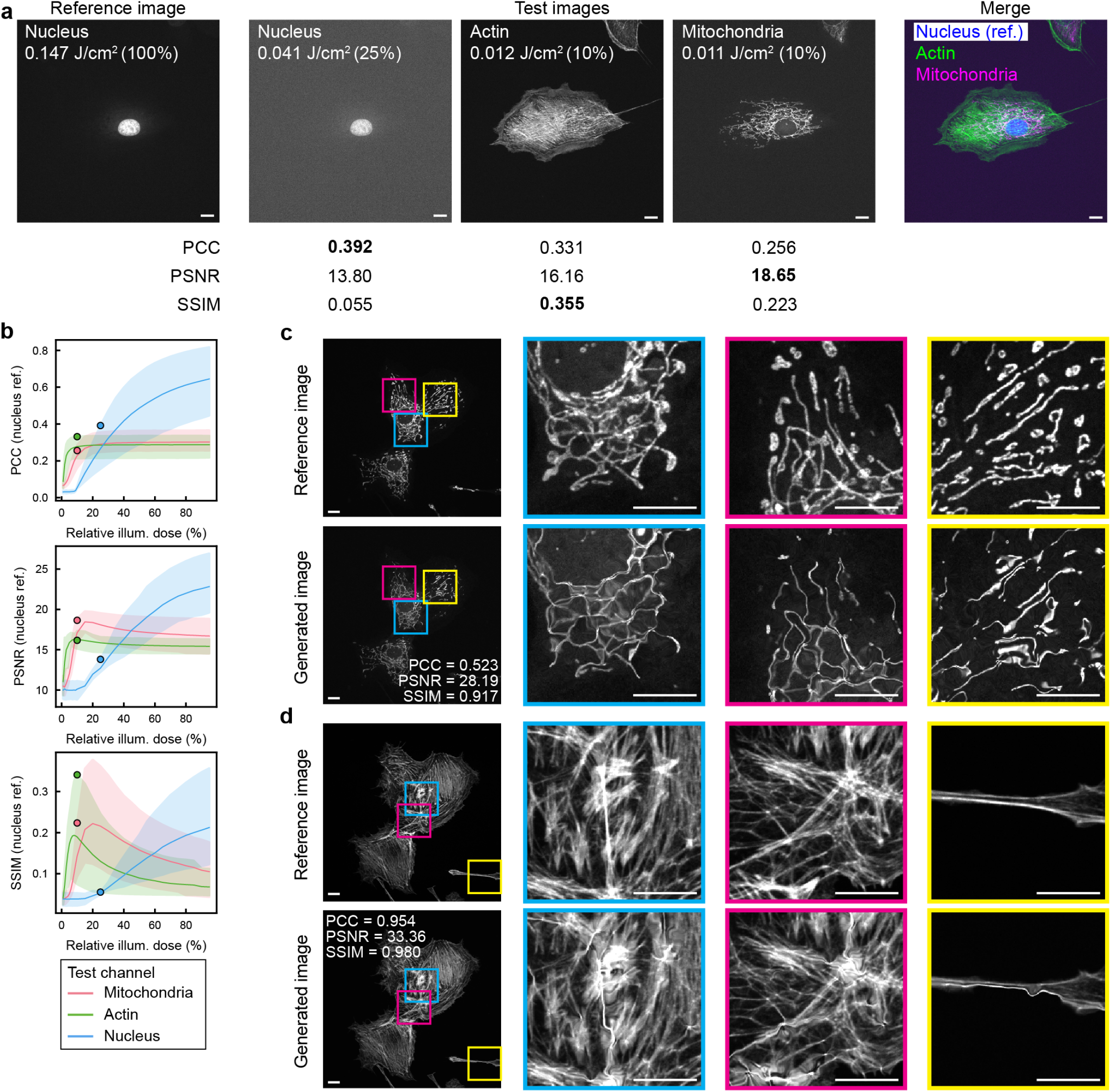
IQM scores can be entirely misrepresentative of biological image content. **a** A high quality image of the nucleus is used as the reference to calculate IQM scores for lower quality images of the nucleus, actin and mitochondria in the same cell. Values in bold indicate the highest score for each metric. **b** Evolution of IQM scores with different illumination doses (here shown as a relative dose given by % of maximum dose for each channel) in three-channel images when using the nucleus channel (100% illumination dose) as the reference. Solid lines indicate mean behaviour over 6 FOVs, with the shaded bands extending to the full range of values. Coloured circles indicate the metric values for the images used in a. **c** An adversarial attack on a mitochondria image (0.07 J/cm^2^) is capable of minimising PCC while maximising SSIM, where the generated image is obviously incorrect. **d** A similar adversarial attack on an actin image (0.08 J/cm^2^) generates an image with high IQM values, but where there are still significant structural defects. All scale bars = 10 µm.

Using an adversarial attack, it was possible to create images with conflicting IQM scores (PCC=0.522, SSIM=0.917, PSNR=28.18; Figure 3c). In this case, the capacity for SSIM and PSNR to report favourably on images that have clearly visible and widespread structural errors raises concern over the use of these metrics for validating image processing methods. Even when the adversarial attack generated images where all three IQMs suggested high quality (PCC=0.952, SSIM=0.894, PSNR=33.00; Figure 3d), noticeable deformities could still be present in images. This underlines that even the PCC – thus far the most robust IQM – can misrepresent finer details of image quality. These results also suggest that for AI-driven image generation and processing tasks, IQMs will not have sufficient structural sensitivity to detect subtle structural hallucinations which are more difficult to perceive than the examples here. Further examples from the adversarial attack can be found in Supplementary Figures S9 and S10.

## Discussion

Here we have demonstrated that commonly used IQMs suffer significant limitations when applied to biological fluorescence microscopy data. While IQMs themselves have been validated in other domains – primarily natural scene image analysis (17) (Supplementary Figure S11) – this is the first in-depth examination of their use in fluorescence microscopy. First, IQMs have little meaning as standalone values. Rather, they offer a relative measure of quality that must be precisely contextualised due to the sensitivity of IQMs to image content. IQMs are only consistent and reliably interpretable in highly similar images, e.g. when comparing quality for different illumination dose versions of the same field-of-view. Even images of the same type of biological structure, labelled identically, and imaged under identical conditions on the same microscope, can produce a broad spectrum of IQM values. Any comparison across studies/benchmarks with different underlying data is at high risk of producing meaningless results. Recently proposed novel IQMs specifically tailored for fluorescence image use (9, 18) are a welcome step towards attempting to tackle the needs of fluorescence-specific image processing, but do not address the challenge of predicting bi-ological analysis performance. Additionally, IQM utilisation is often poorly described in Methods sections, and since numerous factors including image bit depth, normalisation, and the internal hyperparameters of IQMs all alter scores (Supplementary Note 1), increased detail in reporting is vitally needed to make replication and benchmarking credible.

Ultimately, the ability to conduct biological analyses accurately is the central concern in quantitative fluorescence microscopy. Given the increasing development of image processing methods for fluorescence microscopy, it is essential that IQMs should reliably reflect, i.e. be predictive proxies for, downstream biological analysis performance (10, 12), especially when complex transformations are involved (19). Our results indicate that currently used IQMs do not fulfil this requirement. These failings are particularly pernicious where IQMs are used to optimise image processing algorithm development or used as loss functions in deep learning (20), since this may result in the systematic deployment of misguided image processing and analysis tools.

We believe that there is an urgent need for the development of new IQMs suited to fluorescence applications and the task of objectively comparing fluorescence images. In the absence of such ideal metrics, direct quantification of downstream biological analysis performance for tasks of interest should be regarded as the appropriate standard of evidence (3). While such proposals imply significant changes in practice in the field, as well as the need for publicly available datasets to support this, these changes are necessary to ensure that meaningful comparisons can be made between different methods. This will maximise the rigour of methods validation and ultimate impact the new frontiers of research using computation-enhanced fluorescence microscopy.

## Methods

### Imaging samples

#### Schizosaccharomyces pombe cell culture

*S. pombe* was cultured using standard methods(21) on solid (YES agar) and liquid (YES) growth media (Formedium) at a growth temperature of 32°C. Following at least 48 hours of growth, cells in exponential growth were fixed by shaking in 4% paraformaldehyde (Thermo Scientific) for 30 minutes at room temperature. Fixed cells were pelleted, washed and resuspended in 1 ml PBS. For imaging, 35 mm dishes with uncoated polymer coverslips (Ibidi) were first coated with 1 mg ml^−^1 soybean lectin (in water, aliquots stored at - 70°C, Sigma-Aldrich) for 15 min. After washing away the excess lectin with fresh YES, fixed cells were allowed to settle for 30 min in 500 µl PBS. The entire plating volume was replaced with fresh PBS prior to imaging. The strain used expressed sfGFP throughout the cell body (22) (h-ura4+:pact1:sfGFP:terminatortdh1 (AV0890)) (Figures 1, S1).

#### Fixed mammalian cells

Alexa Fluor 488 phalloidin, Mito- Tracker Red CMXRos and DAPI in BPAE cells were imaged in a commercial test slide (FluoCells Prepared Slide #1, Invitrogen) (Figures 2, 3, S5, S8, S9, S10).

#### Fibroblast spheroid culture and preparation

Primary dermal fibroblasts were derived, following collection of normal skin tissue from patients providing informed consent (study approved by the National Research Ethics Service (UK) (14/NS/1073)). Cells were cultured in T25 flasks (Sarstedt) with fibroblast culture media (high glucose DMEM (Gibco) supplemented with 10% FBS (Hyclone), 1% sodium pyruvate (Gibco), 1% penicillin-streptomycin (Sigma) and 4 mM L-Glutamine (Gibco)) and maintained at 37 °C and 5% CO2 until reaching approximately 80-90% confluency. Fibroblasts between passage 8-9 were used to create spheroids using the hanging drop method. After rinsing with PBS (Sigma), cells were incubated for 5-10 minutes at 37 °C with 0.05% Trypsin-EDTA (Sigma), quenched with media once cell detachment occurred and centrifuged at 1200 rpm, 21°C for 5 minutes. Cell pellets were resuspended in media and cells were counted using a haemocytometer (Neubauer Improved) to produce cell suspensions containing approximately 90,000-100,000 fibroblasts. These suspensions were centrifuged at 1200 rpm, 21 °C for 5 minutes and cell pellets were resuspended in 3 ml hanging drop mixture (750 µl media and 2250 µl methocel stock solution (methylcellulose (23)). Approximately 30 µl aliquots of this mixture were pipetted onto the inner lid of a 10 cm square tissue culture disk (Thermo Scientific) and PBS added to the bottom of the dish to prevent dehydration. Dishes were incubated for 24 hours at 37 °C and 5% CO2 to enable the formation of 900- 1000 cell spheroids with a diameter between 200-300 µm.

Collagen gels were created using a liquid collagen solution consisting of 2 mg/ml rat tail-derived collagen type I (Corning) supplemented with 100 µg/ml fibronectin (EMD Millipore Corp.), 2 mM HEPES (Sigma), 14.5 mM NaOH (Sigma), 0.3% w/v NaHCO3 (Sigma) and 10 µg/ml Alexa Fluor 647 NHS Ester (AF647, Invitrogen) made up in Opti-MEM (Gibco), supplemented with 10% v/v FBS. Liquid collagen (prepared on ice) was placed directly onto UV light sterilised 4-well chambered coverslips (ibidi). Rectangular pieces of Mylar plastic film (0.125 mm thickness, Radioshack Pro) were fixed to both ends of individual chamber coverslips with UV curing Norland Optical Adhesive 81 (Norland Products Inc.). Liquid collagen mixture was deposited between the Mylar supports and gels then overlaid with PDMS stamps that were pre-treated with 1% BSA (Roche) made up in PBS. Supports were arranged such that stamping with the PDMS stamps allowed for the formation of micro well-like structures that had a defined distance from the coverslip surface. Gels were left to polymerise at 37 °C in a sterile tissue culture incubator for approximately 2 hours. The PDMS stamps used a SU-8 2100 (MicroChem) master wafer which had 200 µm diameter micropillars at 950 µm pitch, with the stamps and wafer manufactured as described in Phillips *et al*. (24). PDMS stamps cut to dimensions of approximately 1.5 cm by 1 cm were stored in 70% ethanol at 4 °C. Prior to use, stamps were irradiated for 30 minutes under UV light and the micropillars coated with 1% BSA for at least 1 hour. Once polymerised, collagen gels were covered with fibroblast culture media. Spheroids were manually pipetted into the microwells; one spheroid was embedded per gel in a central position and samples left to settle for 15 minutes. Samples were incubated at 37 °C and 5% CO2 for 0-7 days.

In-gel spheroids were fixed in 10% formalin (Sigma) for 2 hours at 37 °C after 24 hours, 48 hours, 72 hours or 7 days incubation, and stored in PBS supplemented with 1% penicillin-streptomycin at 4 °C. At room temperature, cells were permeabilised for 15 minutes with 0.25% (v/v) Triton X-100 made up in PBS (PBST, Sigma) then blocked with 5% BSA solution (in PBST) for 1 hour. Samples were rinsed in PBS, then stained with Alexa Fluor 488 Phalloidin (1:500 dilution, Invitrogen) and Hoechst 33342 (1:250 dilution, Cell Signaling Technology) in 5% BSA overnight at 4 °C, protected from light. Cells were washed in PBS for 30 minutes at room temperature and stored in penicillin-streptomycin supplemented PBS at 4 °C until required for imaging. Although the collagen gel and actin cytoskeleton were fluorescently la-belled, only the Hoechst channel was imaged here.

#### Image acquisition

Images were acquired using a Nikon Eclipse Ti2 inverted microscopey coupled with a Yokogawa CSU-W1 SoRa confocal scanning unit and PVCAM Prime 95B sCMOS camera (Teledyne Photometrics). *S. pombe* and BPAE cell samples were imaged using a 1.49 NA SR HP Apo TIRF 100xAC Oil objective (Nikon), and spheroids were imaged using a 0.75 NA Plan Apo VC 20x DIC N2 air objective (Nikon). Samples were excited using a Lighthub Ultra laser bed (Omicron-Laserage Laserproduckte GmbH) at wavelengths of 405 nm (Hoeschst in spheroids, DAPI in BPAE cells), 488 nm (sfGFP in S. pombe, Alexa Fluor 488 phalloidin in BPAE cells) and 561 nm (MitoTracker Red in BPAE cells). Emitted fluorescence was filtered by a quadband filter (Di01-T405/488/532/647-13x15x0.5, Semrock) and an emission filter prior to detection (447/60 for 405 nm excitation, 525/50 for 488 nm excitation, 641/75 for 561 nm excitation). Images were acquired in 16-bit depth in HDR mode.

Intensity series of increasing illumination doses were acquired from lowest to highest. For *S. pombe*, test images were acquired at the following percentages of maximum laser output: 1%, 2%, 3%, 4%, 5%, 10%, 20%, 30%, 40%, 50%, 60%, 70%, 80%, 90%. Reference images were acquired at 100%, and all images were acquired with 100 ms exposure times. For spheroids, test images were acquired at: 5%, 6%, 7%, 8%, 9%, 10%, 20%, 30%, 40%, 50%, 60%, 70%, 80%, 90%. Reference images were acquired at 100% and all images were acquired at with 100 ms exposure times. For images of actin acquired for filament alignment analysis, test images were acquired at: 1%, 2%, 5%, 10%, 20%, 30%, 40%, 50%, 60%, 70%, 80%, 90%. Reference images were acquired at 100% and all images were acquired with 10 ms exposure times. For three-colour imaging of BPAE cells (Figures 3, S8), test images were acquired for all three colours at: 1%, 2%, 3%, 4%, 5%, 6%, 7%, 8%, 9%, 10%, 15%, 20%, 25%, 30%, 35%, 40%, 45%, 50%, 55%, 60%, 65%, 70%, 75%, 80%, 85%, 90%, 95%. Reference images were acquired at 100% and all images were acquired with 50 ms exposure time. For conversion of percentage maximum output into power, power series at each wavelength were measured after the objective (Thorlabs PM100D) and fitted with a cubic function to produce a calibration curve. Illumination doses in J/cm^2^ were calculated as: *I* = (*P* · *t*)*/A*, where *P* = estimated on-sample power (W), *t* = exposure time (s) and *A* = area of field of view (cm^2^).

#### Synthetic samples

DU145 images used to assess IQM biomarker biases were obtained from Gunawan *et al*., using B-Catenin, DNA, CoxIV, Fibrillarin markers (4). Images used in the localisation classification task were obtained from the Human Protein Atlas (HPA) weakly supervised singlecell classification competition hosted on Kaggle (25). 200 images with a single annotated target were considered, resulting in 16 different classes – Nucleoplasm, Nuclear membrane, Nucleoli, Nucleoli fibrillar center, Nuclear speckles, Nuclear bodies, Endoplasmic reticulum, Golgi apparatus, In-termediate filaments, Actin filaments, Microtubules, Centrosome, Plasma membrane, Mitochondria, Cytosol, Vesicles and punctate cytosolic patterns. Only the green channel, labelled for the protein of interest, was used.

For both sets of images, different levels of signal were simulated in addition to the reference by dividing original intensities by 2,4,8,12,16, or 20 in the HPA data, and by 2 in the DU145s. Poisson noise and Gaussian noise (*σ*=5) was then added through the random package in numpy. Images were then minmax scaled to fill the 8-bit range (Supplementary Note 1).

#### Adversarial image generation

An adversarial attack was performed on the SSIM, PCC and PSNR metrics. The metrics were evaluated in pairs (one pair being in the loss function at a time) with the value of one being maximised and the other minimised. A simple optimisation loop was run, minimising the loss function, which had a component for each of the metrics (one positive and one negative, so one would maximise and the other minimise), and a component for a blurred version of the image to regularise the optimisation, avoiding the optimisation achieving a strong difference in metric values by, for example, changing the background offset or similar relatively uninteresting properties. The weighting of the two metrics within the loss function was adjusted by hand to achieve the best illustration of the potential difference (and to take into account the different scaling of the metrics). The value of PCC was squared to improve the stability of the optimisation, and for pixel-by-pixel optimisation, to avoid having the image simply invert. For the results shown in Supplementary Figure S9 (row 2), the image was optimised on a pixel-by-pixel basis. For the results shown in Figure 3c-d, Supplementary Figure S9 (row 1) and Supplementary Figure S10, an elastic deformation of the image was optimised.

### Image processing

#### Mean filtering

Mean filtering was performed using the `uniform_filter` function in the ndimage package in scipy (26), with a patch size of 5. All other parameters were left at the default values.

#### Non-local means filtering

Non-local means filtering was performed using the `denoise_nl_means` function in the restoration package in scikit-image(27). For each image, sigma was estimated using the `estimate_sigma` function in scikit-image.restoration, and consistently a patch size of 5 and a patch distance of 11 was used.

#### Pre-trained AI

The pre-trained AI denoising used here was the Nikon Denoise.AI method (14), which aims to remove shot noise from confocal images. This is a pre-trained convolution neural network provided as a plugin to the NIS Elements acquisition and analysis software. Neither the network architecture nor the training data are publicly available; the only information about the training data is that it was trained on “several thousand examples of acquired Nikon confocal data”. For all acquired images, Denoise.AI was applied online during acquisition, with denoised acquisitions interleaved with raw acquisitions in the power series (NIS Elements AR 5.42.06). For synthetic noisy images, Denoise.AI was batch applied offline following image conversion of 8-bit png to 8-bit tiff files (NIS Elements AR 6.02.03).

#### Custom AI

The AI image restoration network used here was CARE (15) (content aware image restoration). CARE learns how to restore images via providing training data comprising pairs of the same field of view imaged at high and low qualities. Models were trained using the CAREamics python library (28). All models were trained using MAE loss for 100 epochs with a batch size of 4 and a 0.8 validation split. For real images, a patch size of 128 × 128 was used, and for synthetic images a patch size of 64 × 64 was used.

For all datasets, two models were trained per illumination dose (or noise level, for synthetic data). At each illumination dose, fields of view were split randomly into two groups (along with the maximum illumination dose fields of view, which were used as the ground truth). Each group was augmented with flips and rotations, and then a CARE model trained for each group. The CARE model from the first group of images was used to make predictions for the second group of images, and vice versa, ensuring independence of training data from test data. The training data sizes were: S. pombe – 30 1412 × 1412 pixel images, actin – 30 1412 × 1412 pixel images, spheroid – 33 1412 × 1412 pixel images, HPA synthetic data – 1600 256 × 256 pixel images. For the HPA dataset, the CARE models at each noise level included all labels.

#### Image Quality Metrics

Peak signal noise ratio (PSNR) is designed to measure the ratio between signal strength and noise. Structural similarity index (SSIM) is inspired by human perception consisting of terms to accounts for luminance, contrast and structure. Both PSNR and SSIM were calculated using the metrics package in skimage (version 0.24.0). SSIM calculations in Figures 1d and 2e were performed using default parameters; all other SSIM calculations were performed to match performance to the original SSIM paper implementation (`gaussian_weights=`True, `sigma=`1.5, `use_sample_covariance=`False), with SSIM parameterisation described further in Supp Note 1. Pearson’s correlation coefficient (PCC) is statistically motivated, modelling correspondence of pixels as is done for variables across many domains. Data in Figures 1d, 2e and 3c-d were min-max normalised prior to IQM calculation, and all other data were percentile normalised prior to IQM calculation. Supplementary Note 1, Tables S2 and S3, and Figure S11 contain a full description of the metrics along with their default parameters and the effect of different normalisation strategies.

### Downstream analysis

#### StarDist nucleus segmentation

StarDist model training and predictions were made using the python implementation of StarDist (version 0.4.1). To ensure that segmentation performance only depended on the properties of the images being segmented, separate StarDist models were trained for each different image type (raw, mean-filtered, NLM processed, pre-trained AI denoising, custom AI denoising) with a separate model for each illumination dose within that image type. StarDist training was performed using the default suggested parameters, with 32 rays per polygon and 200 training epochs. The training data was composed of 76 manually annotated 256 × 256 pixel patches from 24- and 48-hour post-seeding spheroids to ensure that training data contained a mix of high-density and low-density regions; this corresponded to a total of 3000 annotated nuclei in the training data. The raw training data was imaged at the same illumination doses and processed in the same way as the test data, and the dataset was augmented using random flips and rotations prior to training. 15% of the training data was used for validation each time the model was trained.

Nucleus segmentation accuracy was quantified using the aggregated Jaccard index (AJI) as described in Kumar et al. (29). This method accounts for the segmentation of each matching nucleus (true positive) between the ground truth and test image, as well as false positives and false negatives in the test image. In all cases, StarDist segmentation of the reference image was used as the ground truth to assess the accuracy of StarDist segmentations of raw and processed test images.

#### Alignment by Fourier Transform (AFT) analysis

AFT analysis of actin images was performed using a patch size of 80 pixels with an overlap of 50%, resulting in 33 × 33 blocks per image (AFT release version ‘Code for [Marcotti et al. 2021]’ on GitHub) (30). No local filtering or masking was applied to the images during AFT analysis. The angle maps produced by AFT were saved, and Pearson’s Correlation Coefficient was only calculated between test and reference AFT angle maps for blocks where the reference fluorescence intensity was above the Otsu threshold.

#### Human Protein Atlas classification

HPA classification was performed using ResNet18 (31), a representative convolutional neural network architecture commonly used for image classification. The model was trained for 100 epochs from scratch, optimised using stochastic gradient descent (learning rate = 0.0001, momentum = 0.9), using a fully connected final layer with 16 output features corresponding to the 16 protein sub-localisation classes, activated with the log softmax function. Experiments were repeated using 5-fold stratified cross validation.

## Data availability

All raw and denoised image data used in this study is available on the BioImage Archive, including training data for StarDist models and StarDist predictions across all illumination doses and denoising modalities.

## ACKNOWLEDGEMENTS

We thank the Nikon Imaging Centre at King’s College London, particularly Dylan Herzog, for assistance with and maintenance of spinning disk microscopy equipment. We thank Gautam Dey (EMBL, Heidelberg) for *S. pombe* cells. We thank Tanya J. Shaw from King’s College London for providing access to the primary dermal fibroblast cells used. We thank Michelangelo Colavita from Nikon UK for helping develop scripts for power series image acquisition with online denoising, and Nikon UK for the loan of a spinning disk microscopy. We thank Thomas A. Roberts for providing the photograph in Supplementary Figure S11, and Ripley, Artemis and Scamp for posing in it. IG is supported by the Australian Government Research Training Program (RTP) Scholarship. NA is supported by a Mechanics of Life Leverhulme Doctoral Scholarship. RJM and SCo are supported by UK BBSRC grant BB/X01858X/1. JGL is supported by a University of New South Wales Scientia Research Fellowship, a Ramaciotti Biomedical Research Award, an ARC Development Project grant (DP170103599), NHMRC Ideas Grants (GNT1184009, GNT2012848, GNT2028506), and a Tour de Cure Pioneering Grant (RSP-547-FY2023). SCu and RJM are supported by funding from Royal Society University Research Fellowship URF\R1\211329.

## AUTHOR CONTRIBUTIONS

**Ihuan Gunawan**: conceptualization, formal analysis, investigation, methodology, software, visualization, writing – original draft (lead), writing – review and editing. **Richard Marsh**: conceptualization, methodology, writing - review and editing. **Nandini Aggarwal**: methodology, resources. **Erik Meijering**: conceptualization, supervision, writing - review and editing. **Susan Cox**: conceptualization, formal analysis, software, visualization, writing – review and editing. **John G Lock**: conceptualization, methodology, resources, supervision, writing – review and editing. **Siân Culley**: conceptualization, data curation, formal analysis, funding acquisition, investigation, methodology, resources, software, visualization, writing – original draft, writing – review and editing.

## COMPETING FINANCIAL INTERESTS

The authors declare no competing financial interests.

## Supplementary Information

**Fig. S1.**
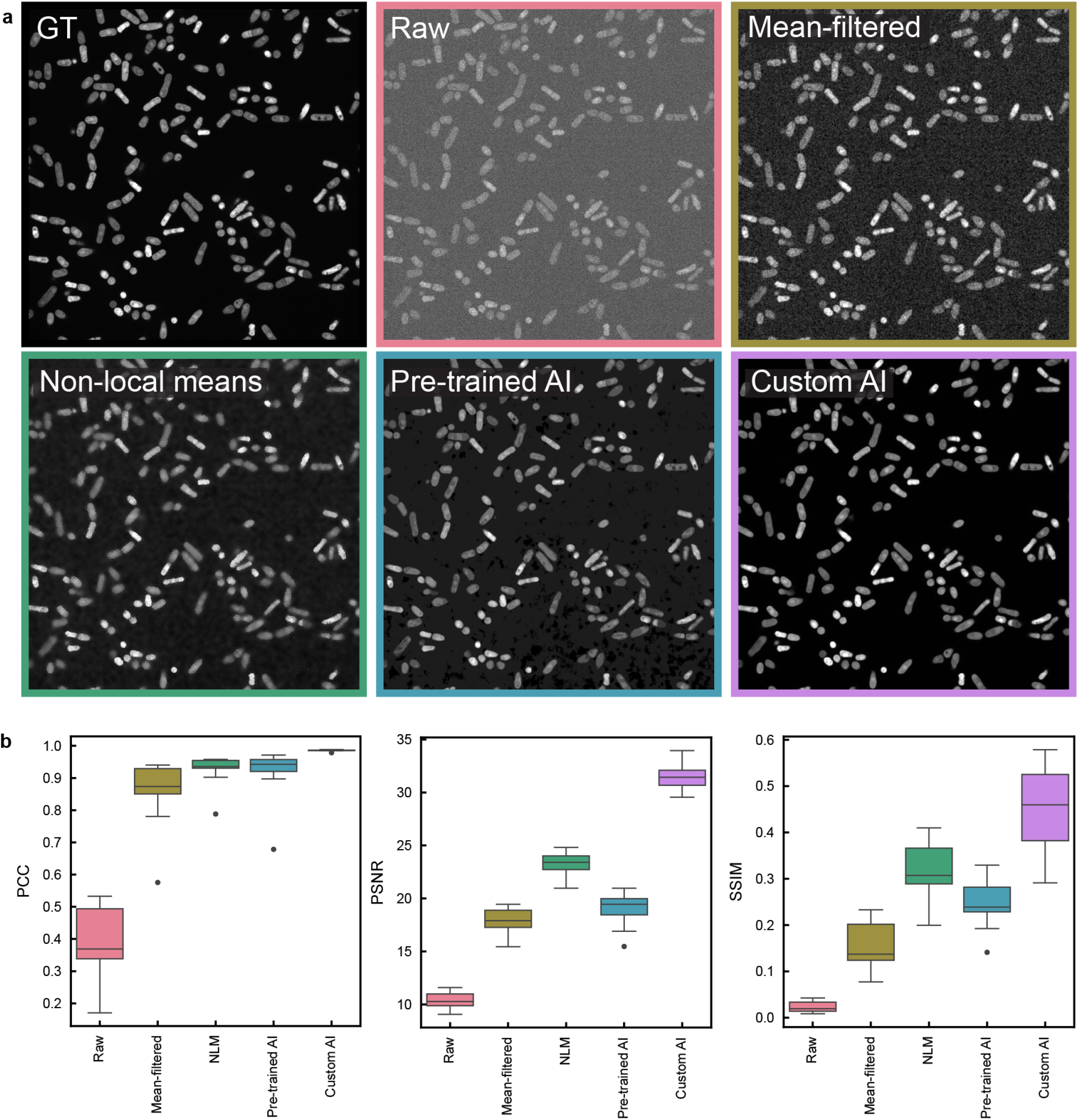
Typical application of IQMs to image processing in microscopy. **a** Representative images of the same field of view for *S. pombe* expressing sfGFP. Ground truth (‘GT’) image is acquired at a high illumination dose (0.76 J/cm^2^), ‘Raw’ image is acquired at low illumination dose (0.02 J/cm^2^), and the results of four image processing methods applied to the Raw image are displayed. **b** Box plots showing bulk IQM values for 21 fields of view acquired at 0.02 J/cm^2^ before (Raw) and after processing. Boxes show three quartile values of distribution, whiskers extend to 1.5 interquartile range. Values outside this range are plotted as individual points.

**Fig. S2.**
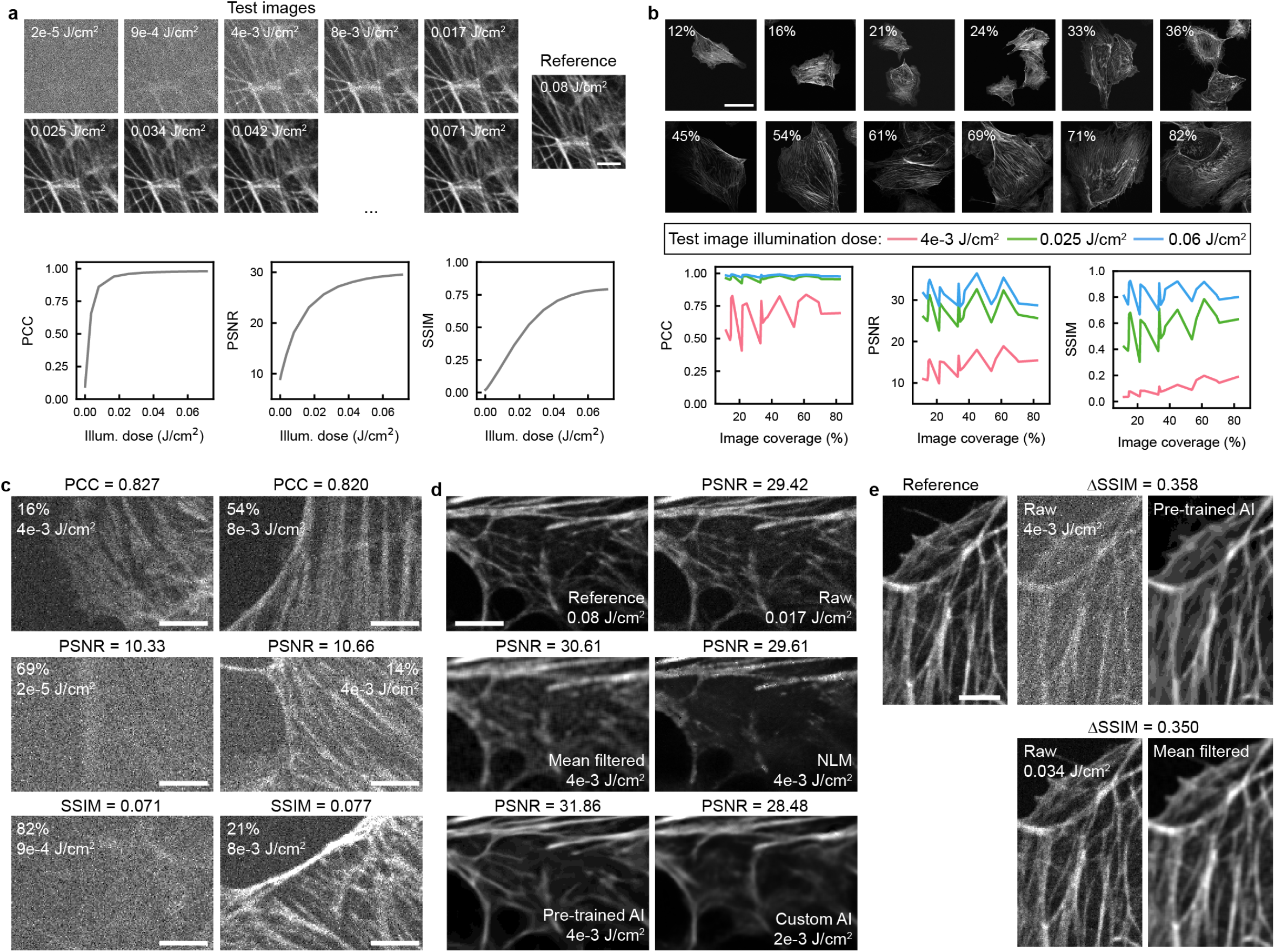
IQM behaviour on raw and processed microscopy images of fluorescently-stained actin. **a** Top: Crops from a series of images of fixed mammalian cells stained with phalloidin-488. Bottom: IQM values for one whole field of view as a function of illumination dose. **b** Top: Whole fields of view of fluorescently-labelled actin containing different image coverages, acquired at 0.08 J/cm^2^. Bottom: IQM values plotted as a function of image coverage, for three different test illumination doses. **c** Crops from images where the whole fields of view have a similar IQM value (top: PCC, middle: PSNR, bottom: SSIM) despite acquisition with different illumination doses. **d** Crops from images either raw or following image processing, where each whole field of view has a PSNR ∼ 30. **e** Crops from images before and after image processing, where for both images processing has improved the SSIM score by ∼ 0.35. In the top panel this corresponds to a substantial improvement in visual quality, whereas in the bottom panel similar features are visible both beofre and after processing. Scale bars: a - 5 µm, b - 50 µm, c - 5 µm, d - 5 µm, e - 5 µm.

**Fig. S3.**
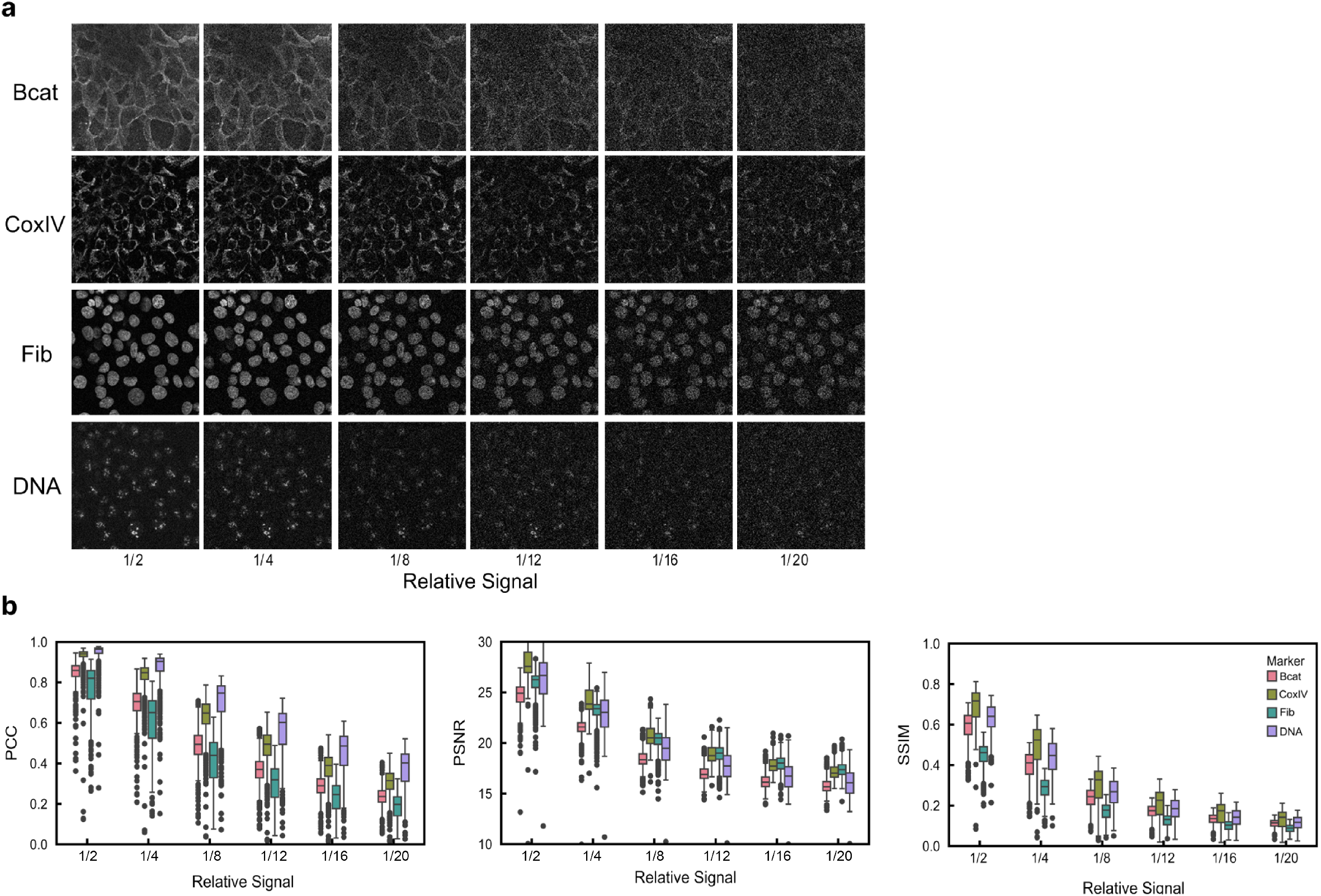
Bulk IQM assessment of images for various markers. **a** Example of image fields for each mark with varying signals (relatively scaled as labelled). **b** Box plots showing range of resulting IQM scores per marker. A clear downward trend observed with less signal. Each box represents scores calculated from 583 different fields. Boxes show three quartile values of distribution, whiskers extend to 1.5 IQR. Values outside this range are plotted as individual points.

**Fig. S4.**
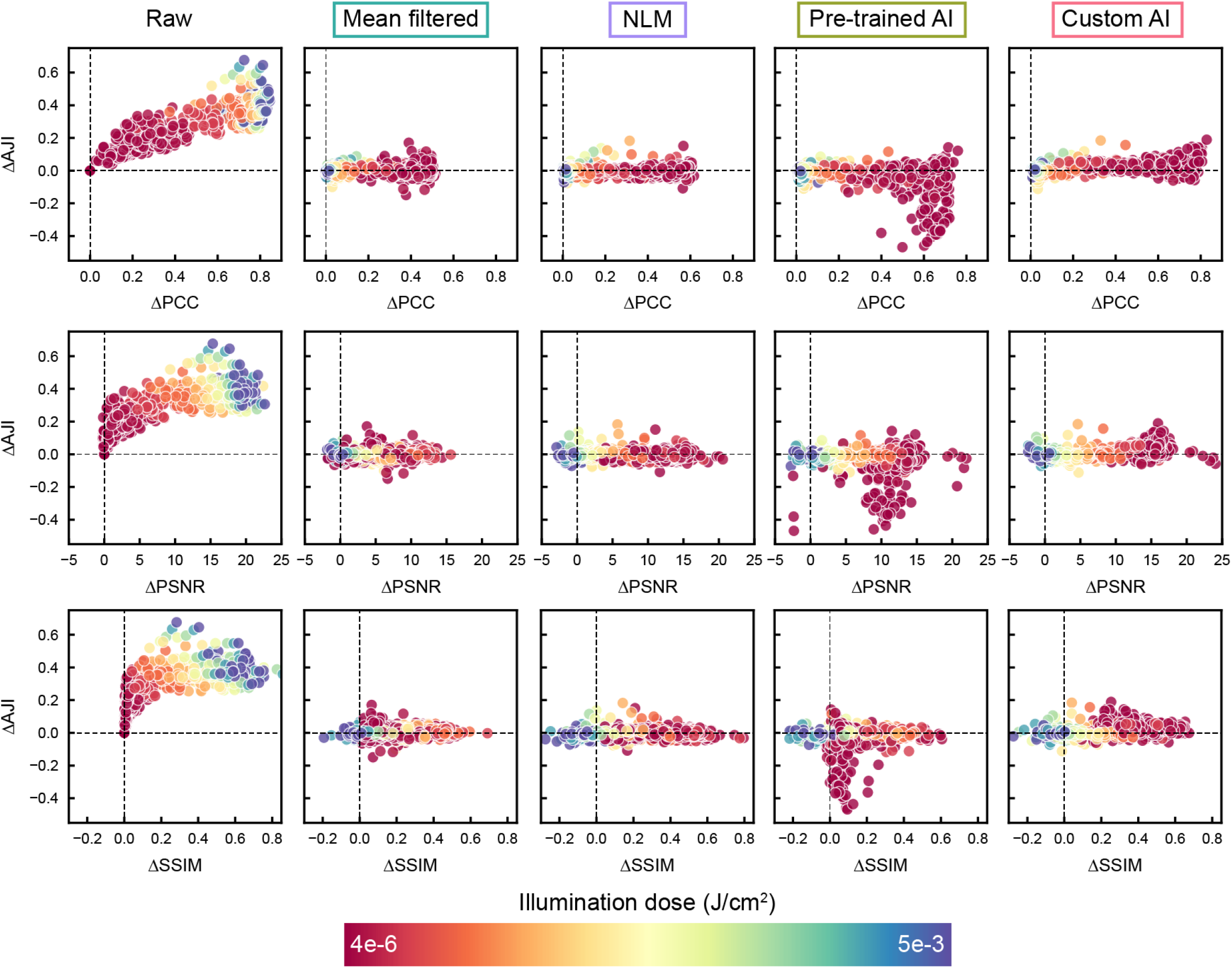
Impact of illumination dose on denoising performance and spheroid nucleus segmentation accuracy. Grouped data from Figure 2c and Figure S7a are split into data points from individual images and colour-coded according to the illumination dose used to acquire the raw image. For the Raw data, ΔAJI and ΔIQM values are calculated relative to the values from the lowest illumination dose for each field of view. For the denoised data, ΔAJI and ΔIQM values are calculated as the difference between the denoised and raw AJI and IQM values for each illumination dose.

**Fig. S5.**
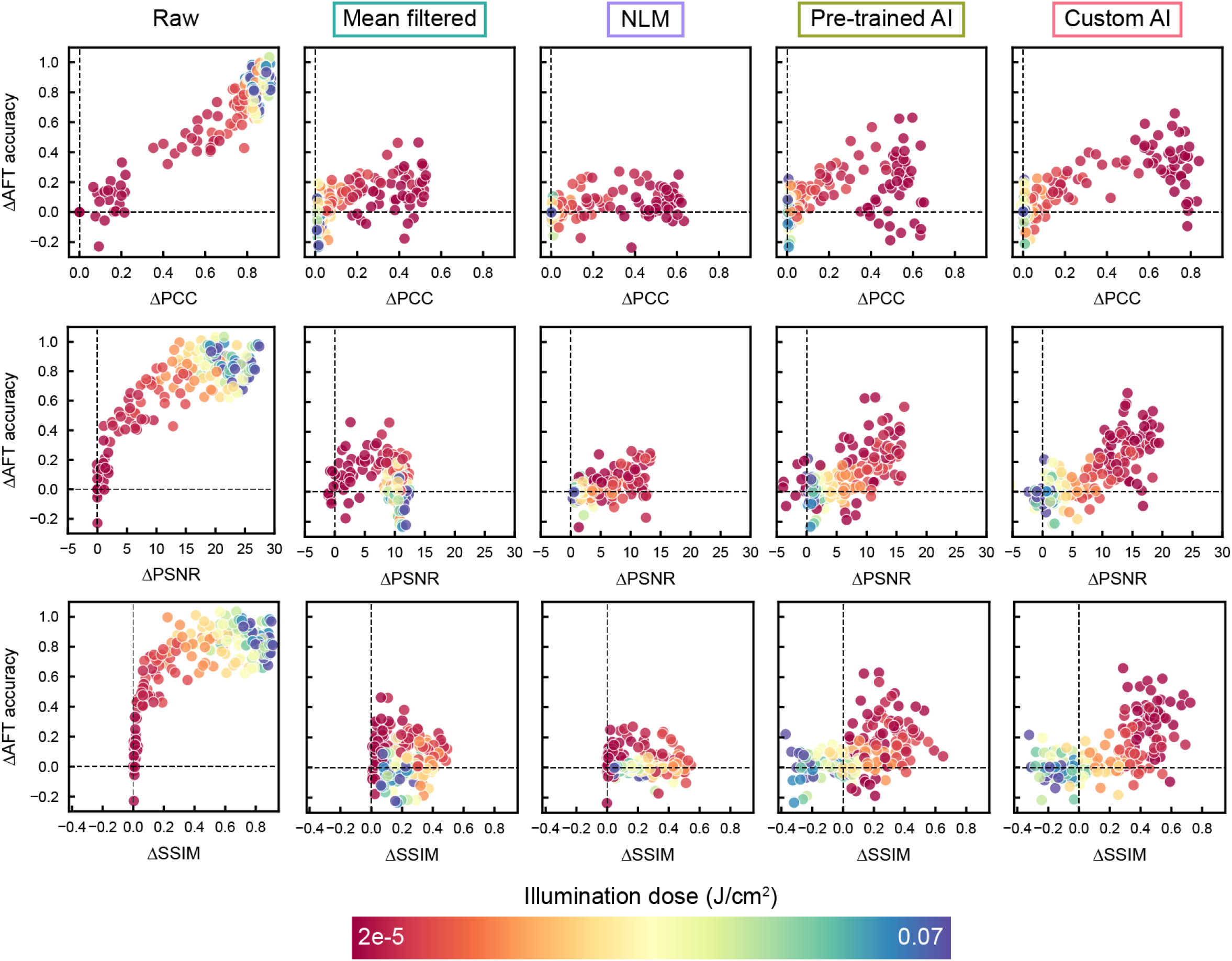
Impact of illumination dose on denoising performance and actin filament orientation analysis accuracy. Grouped data from Figure 2d and Figure S7b are split into data points from individual images and colour-coded according to the illumination dose used to acquire the raw image. For the Raw data, ΔAJI and ΔIQM values are calculated relative to the values from the lowest illumination dose for each field of view. For the denoised data, ΔAJI and ΔIQM values are calculated as the difference between the denoised and raw AJI and IQM values for each illumination dose.

**Fig. S6.**
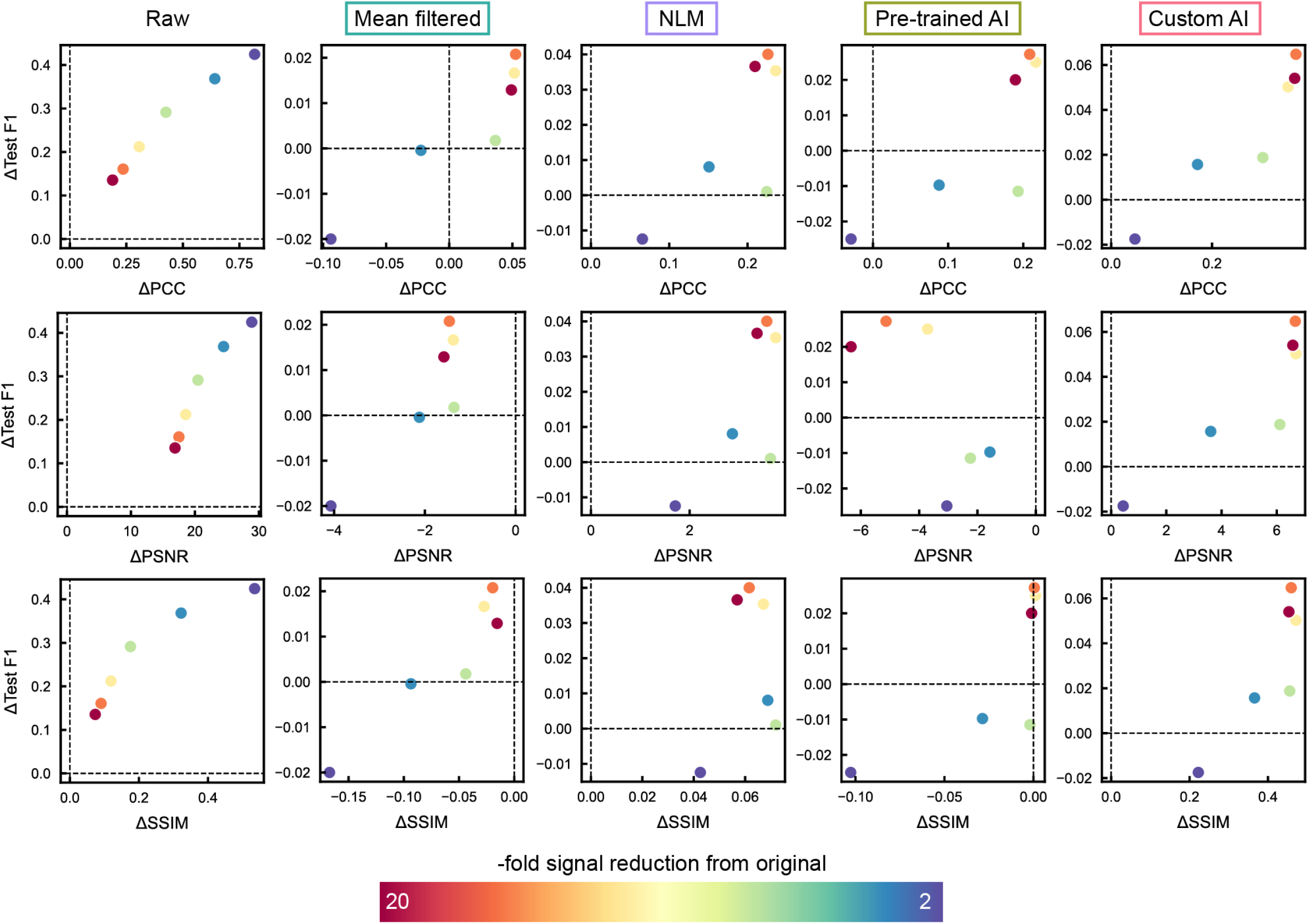
Impact of synthetic signal reduction on denoising performance and whole image classification accuracy. Grouped data from Figure 2e and Figure S7c are colour coded according to the strength of signal reduction applied to generate synthetic noisy images.

**Fig. S7.**
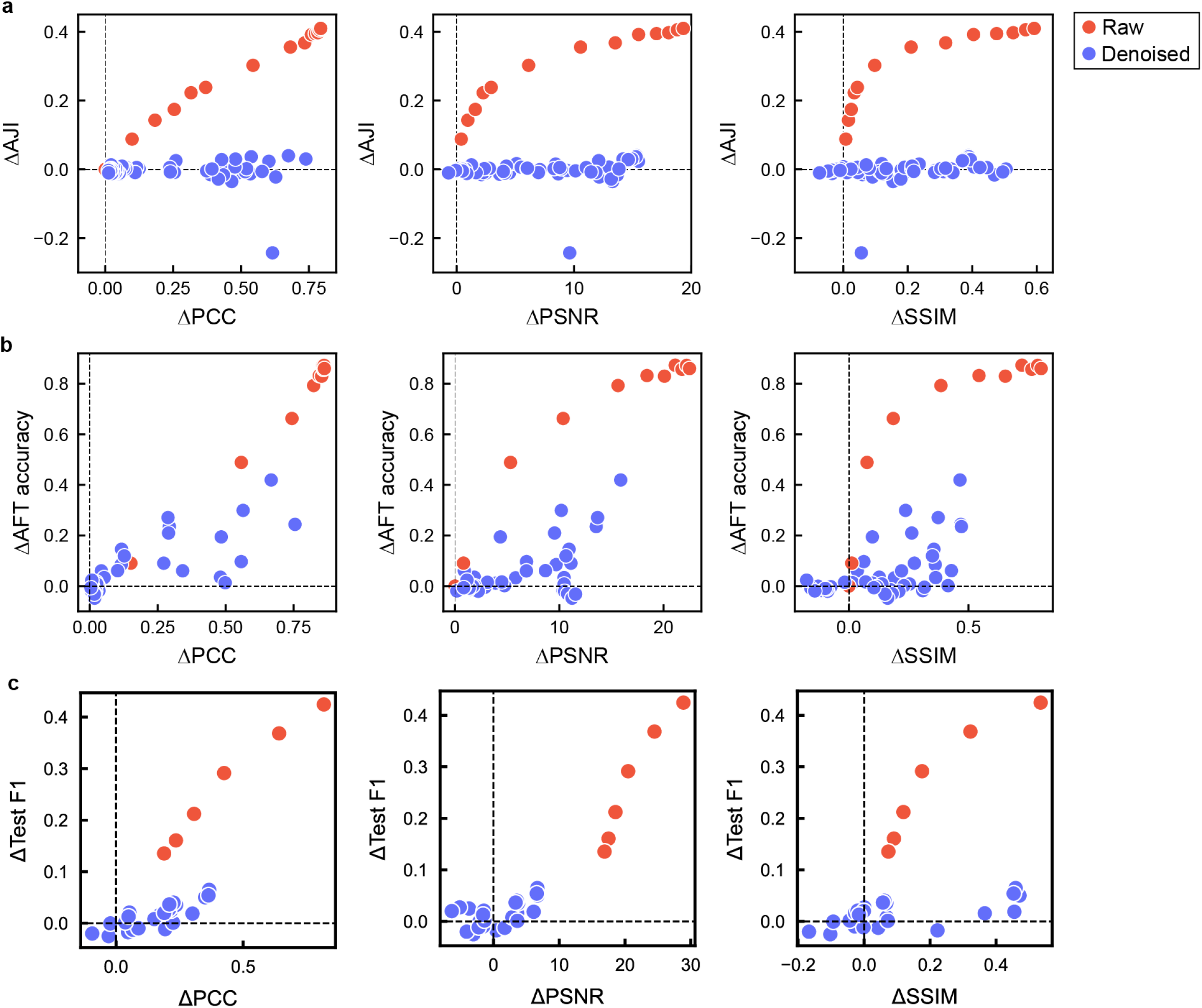
Comparison of image analysis accuracy on denoised and raw images. **a** Plots of segmentation accuracy (ΔAJI) against ΔIQM for spheroid data. Denoised data are the same as in Figure 2c, but all methods are coloured blue here for clarity. Changes in AJI and IQM value for the raw images are comparisons against the lowest illumination dose image for each illumination dose acquired. **b** As in **a**, but for filament alignment accuracy (ΔAFT). **c** As in **a, b** but for whole image classification accuracy (ΔTest F1). Here, changes in F1 score and IQM value are comparisons against classification performance of reference images (no simulated noise).

**Fig. S8.**
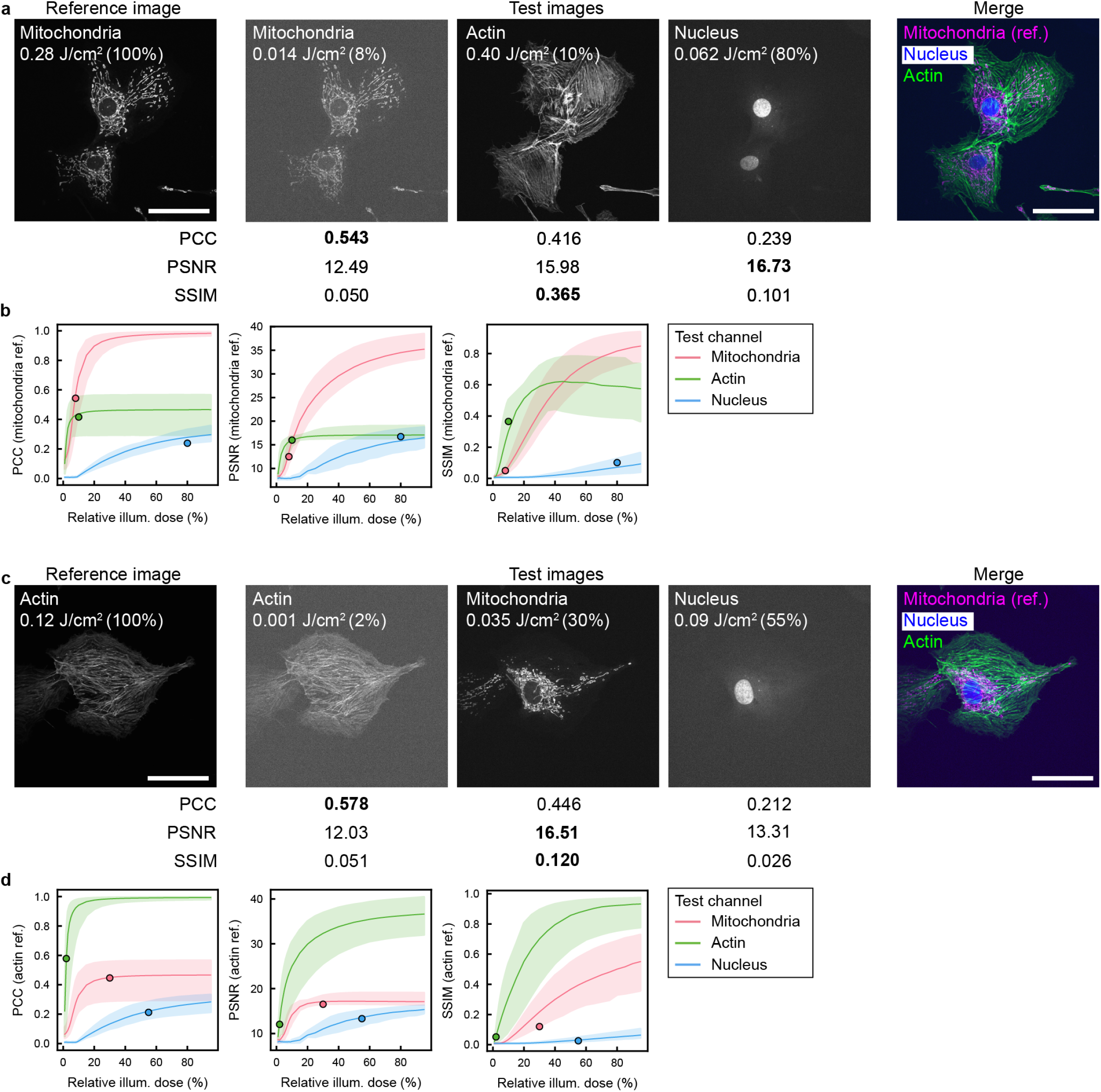
Further examples of IQMs ‘favouring’ incorrect biological structures. **a** Images of mitochondria, actin and nucleus stains in the same field of view. The reference image for the IQMs is the mitochondria image at high illumination dose, and the test images are a low illumination dose mitochondria image, medium illumination dose actin image, and high illumination dose nucleus image. Quality metrics are calculated for each image against the reference mitochondria image; numbers in bold indicate the image with the highest scoring IQM. **b** Plots showing the relationship between relative test image illumination dose and IQM for each channel, where in each case the reference is the highest illumination dose mitochondria image for each field of view. Points indicate the images in **a. c** As in **a**, but with a high illumination dose actin image as the reference, and low illumination dose actin image, high illumination dose mitochondria image, and high illumination dose nucleus image as the test images. **d** As in **b**, but with the highest illumination dose actin images as the reference, and points corresponding to the field of view in **c**. Scale bars = 50 µm. In all graphs, solid line is mean of N=6 fields of view, and shaded areas extend from minimum to maximum.

**Fig. S9.**
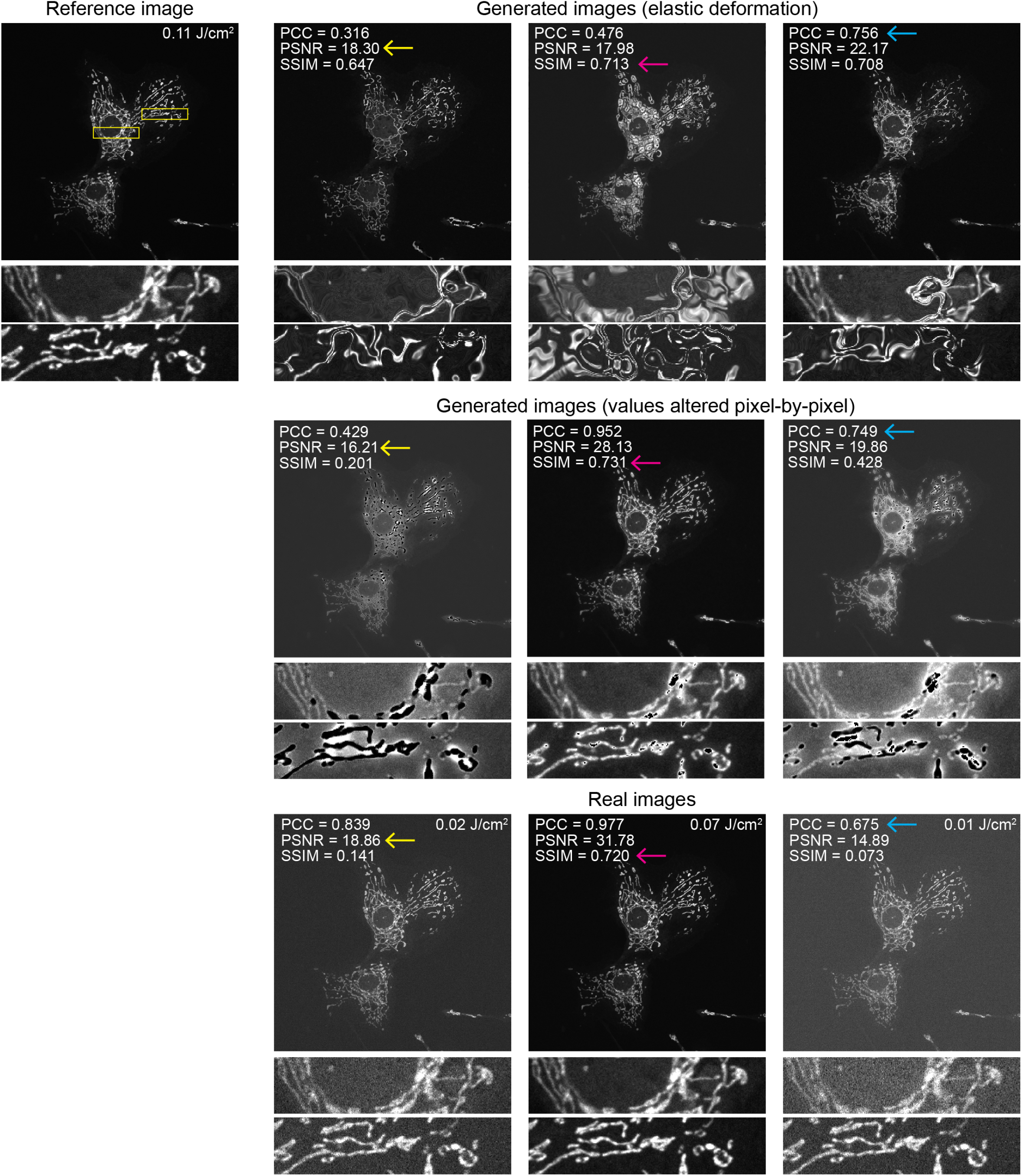
Examples of adversarial images obtained via two different image generation strategies and their IQM values compared to real images. An adversarial attack was performed on various combinations of PCC, SSIM and PSNR metrics (see Figure S10) by optimising an elastic deformation (top row) or pixel-by-pixel value alteration (middle row) of a 0.07 J/cm^2^ image of mitochondria. IQMs were calculated on 1-99.8% normalised images between the reference image and the generated images. The bottom row shows examples of real acquired images with similar IQM values to the generated images for context (yellow arrows = similar PSNR, magenta arrows = similar SSIM, cyan arrows = similar PCC).

**Fig. S10.**
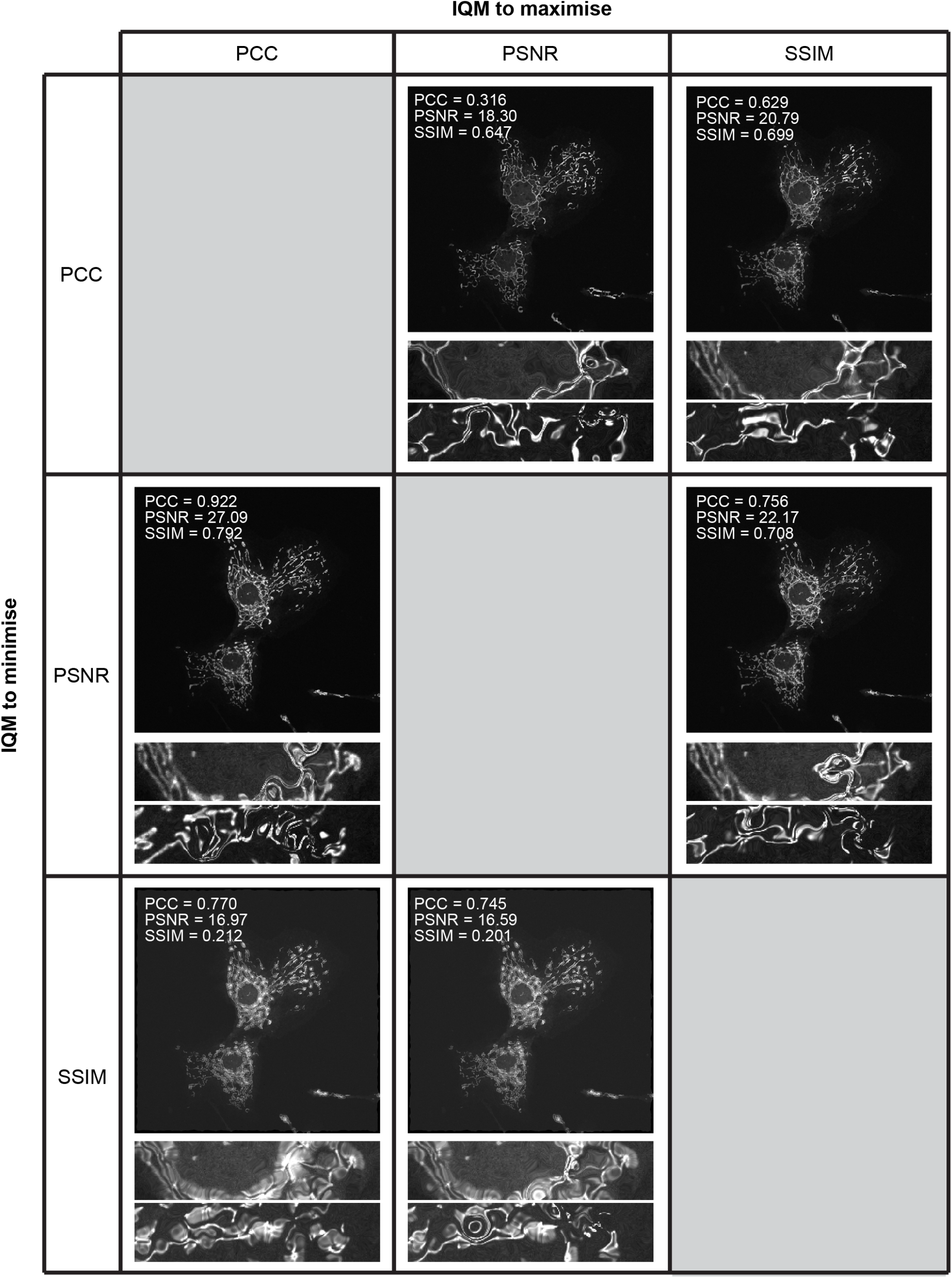
Generated images from difference adversarial attack strategies with elastic deformation. Adversarial attacks were performed on the same starting image (mitochondria imaged at 0.07 J/cm2 illumination dose). Columns indicate the IQM which the attack tried to maximise, and rows indicate the IQM which the attack tried to minimise. IQMs were calculated on 1-99.8% normalised images using the 0.11 J/cm^2^ mitochondria image as the reference (see Figure S9). Zoomed insets are from the same region as shown in Figure S9.

**Table S1.**
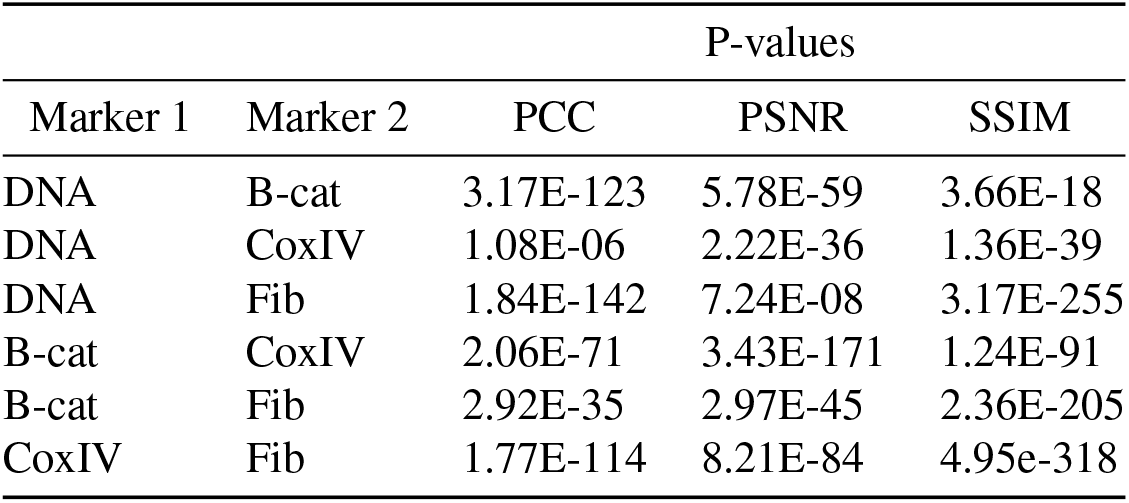
Significance testing for IQM ranges for different subcellular markers. P-values are shown for testing the significance of differences between IQM values reported between pairs of markers.

## Supplementary Note 1: Image quality metric formulae, parameters and implementations

### A. Peak signal-to-noise ratio (PSNR)

The peak signal-to-noise ratio is defined as:

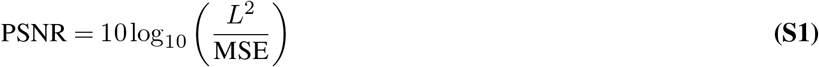

where *L* is the dynamic range of the image (discussed further below) and MSE is the mean-squared error between the two images, defined as:

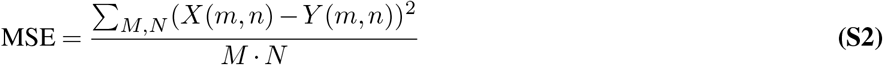

where *X* and *Y* are the reference and test images respectively, each having width *M* pixels and height *N* pixels. PSNR originates in signal processing and has been extended to images; there is no reference for PSNR as an image quality metric, however common implementations (for example scikit-image and MATLAB’s Computer Vision Toolbox) do not differ in their formulae.

### B. Structural Similarity Index Measure (SSIM)

The structural similarity index measure is defined as (1):

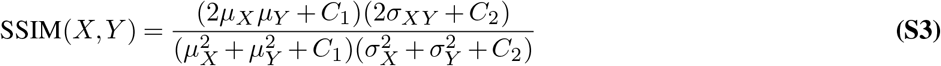

where *X* and *Y* are the reference and test images as for the PSNR above. *µ*_*X*_ and *µ*_*Y*_ are the mean intensities of the reference and test images, 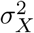 and 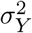 are the variances of the reference and test images, and *σ*_*XY*_ is the covariance of the reference and test images. The constants *C*_1_ and *C*_2_ are present to prevent instabilities when 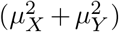 or 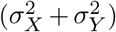 are close to zero, and are defined as:

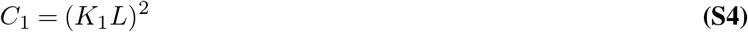

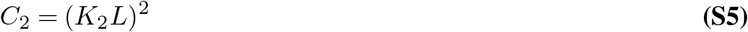

where *L* is the dynamic range of the image. Common implementations set the default values of *K*_1_ and *K*_2_ and 0.01 and 0.03 respectively, as per the original paper. The original paper notes that these values are somewhat arbitrary. The original rationale for developing the SSIM metric was to combine structural, contrast and luminance into a single metric which reproduces key features of the human visual system. Further, the metric was only originally tested on natural scene images and not any biomedical images. Studies in magnetic resonance imaging, for example, have since highlighted discrepancies in SSIM performance and subjective image quality assessment by radiologists (2). Additional parameters are provided in software implementations which are not captured by the SSIM formula. For example, the scikit-image implementation has the option to calculate the SSIM in a window-wise fashion across the image, with further option to spatially normalise the mean and variance of the values within each window by a normalised Gaussian kernel. The MATLAB implementation has the option to change the weighting by which the structure, luminance and contrast terms are weighted within the SSIM formula (presented above with all weights equal to 1). As MATLAB has an option to output an SSIM map, we assume that it also calculates SSIM patchwise across the image, although this is not described in the documentation.

Some microscopy papers use an extension of SSIM, multi-scale (MS)-SSIM (3) as a quality metric. This calculates the luminance, contrast and structure terms of the SSIM metric between the test and reference images with progressive downsampling of the images (typically following Gaussian blurring of the images), and then combines these exponential weights:

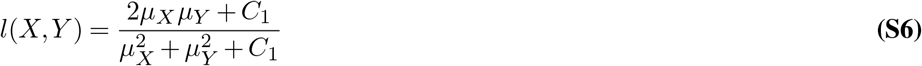

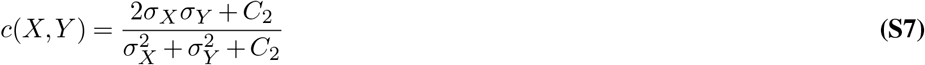

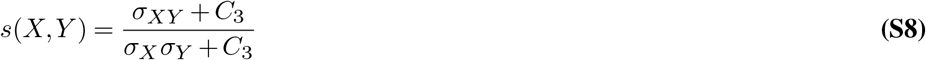

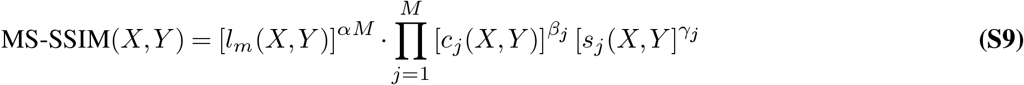

where *l, c* and *s* are the luminance, contrast and structure terms respectively. *C*_1_ and *C*_2_ are defined as above, and *C*_3_ = *C*_2_*/*2. The downsampling scales are represented by *j*, and range from 1 to *M*. *α, β* and *γ* are used to adjust the relative contributions of the three terms, and per the original paper parameters are selected such that *α*_*j*_ = *β*_*j*_ = *γ*_*j*_ for all values of *j*. Furthermore, to normalise cross-scale settings, 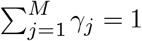. The values of parameters were determined in the original paper by asking 8 subjects to compare the quality of 10 sets of natural scene images presented with different degrees of distortion across 5 scales. The obtained parameter values were: *β*_1_ = *γ*_1_ = 0.0448, *β*_2_ = *γ*_2_ = 0.2856, *β*_3_ = *γ*_3_ = 0.2363 and *α* = *β*_1_ = *γ*_1_ = 0.1333. The authors note that this parameter selection is somewhat crude and that the parameters themselves are “abstract”. It should be underlined that the entire rationale of the MS-SSIM metric is, as for SSIM, to mimic the processing power of the human visual system on natural scene images, which is not typically how microscopy images should be assessed.

There is no MS-SSIM implementation provided in the scikit-image python package. Both the pytorch-metrics package and MATLAB have an MS-SSIM implementation where the *k*_1_, *k*_2_, *α, β* and *γ* parameters are set to the values described in the original paper. Both implementations also provide the opportunity to change the number of scales (*M*, with a default of 5), and the sigma of the Gaussian function used to blur the images prior to downsampling (default 1.5 pixels). pytorch-metrics also has an option for normalisation of the MS-SSIM metric, but this is only intended for use when MS-SSIM is being applied as a loss function. When independently demonstrated for use in microscopy, the same parameter values are used as for the original paper and package defaults (4).

### C. Pearson’s correlation coefficient

Pearson’s correlation coefficient is calculated as:

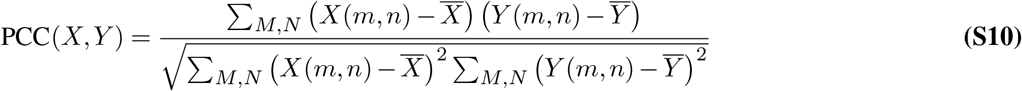

There are no parameters associated with Pearson’s correlation coefficient, and it is provided as a built-in function in most major numerical packages (for example, the `pearson_corr_coeff` function in scikit-image.measure and the `corr` function in MATLAB).

### D. Image dynamic range and normalization

The calculation of both the PSNR and SSIM metrics depend on the dynamic range of the image, denoted by *L* in Equations S1, S4, S5. Natural scene images are usually 8-bit, meaning a maximum dynamic range of 255 (2^8^ − 1). For many natural scene images, most of the bits are represented in the image histogram (Figure S11a, blue histograms). However, in fluorescence microscopy, raw data from the microscopy is typically 8-bit (*L* = 255), 12-bit (*L* = 4095) or 16-bit (*L* = 65535). For higher bit depths, it is very uncommon for all bits to be represented in the image histogram; particularly for 16-bit camera-based imaging, most pixels have intensities (represented by bits) ≤ 10^4^, with higher bits not represented in the image histogram (Figure S11b, blue histograms). Furthermore, image processing operations frequently produce images with double or floating point precision, for which the dynamic range is not well-defined.

While it may seem trivial, the dependence of PSNR and SSIM on dynamic range means that allowing default selection of parameters in software packages can give unexpectedly high metric values (Table S2). For natural scene images, where the bit depth is well-filled, both the PSNR and SSIM values calculated on the raw data (*L* = 255) correspond intuitively with perceptual values in image quality. However, for microscopy images, PSNR and SSIM values calculated on the raw data (*L* = 65535) are higher than one might expect from visual inspection of the data. Particularly for the SSIM, calculation with the ‘native’ data range of the raw images gives a very high value for an image where the details are seen to be noisy and difficult to resolve. This is not unexpected behaviour if we look at the formulae for PSNR and SSIM; the pixel values in the high-quality image are 79-2089, and the pixel values in the reference image are 80-2751. While there are noticeable differences between these pixel values, these are orders of magnitude away from the contribution of *L* in the PSNR and SSIM formulae. As a result, *L* dominates the MSE term in the PSNR, and the *C*_1,2_ coefficients dominate the mean and variance terms in the PSNR.

For microscopy data, we recommend that any calculation of the PSNR and SSIM should be performed on images that have been converted to floating point precision and normalised from 0-1 (i.e. such that *L* = 1); this also removes any problems that may arise if there are differences in bit depth between raw and processed images. The two normalisation strategies used in this paper are min-max normalisastion and percentile normalisation. For the raw and processed images of *S. pombe*, actin cytoskeleton, fibroblast nuclei, 3-colour stained cells, and adversarial generated images in Figures S9, S10, percentile normalisation was used as these images were all acquired using an sCMOS camera (or in the case of the generated images, were to be compared with images acquired using an sCMOS camera), which are known to have out-of-distribution hot and cold pixels. This was performed as:

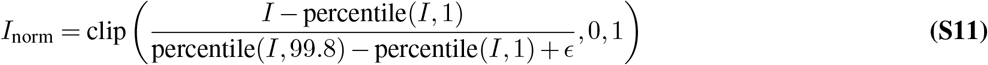

where *I*_norm_ is the percentile normalised image, *I* is the original image, percentile(*I, p*) is the *p*th percentile (pixel value where *p*% of the pixels have lower intensity) as calculated using the `percentile` function in numpy, *ϵ* is a small constant (1*e* 20) to prevent division by zero, and clip denoised clipping of the pixel range to the range [0, 1] to ensure consistency with *L* = 1. For the DU145 and HPA images, which are laser-scenning confocal microscopy data, out-of-distribution pixels should not be produced by the detector and so min-max normalisation was performed as follows:

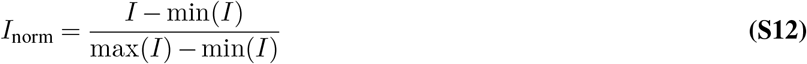

where *I*_norm_ and *I* are as above, min(*I*) is the minimum pixel value in *I*, and max(*I*) is the maximum pixel value in *I*. Normalising the microscopy images in Figure S11 prior to calculating IQMs has a significant impact on the PSNR and SSIM values, particularly for the lower quality image (Table S2). In contrast, normalising the 8-bit natural scene images has a less marked effect on IQM values, and percentile normalisation probably has a deleterious effect on metric calculation due to the absence of outlier pixel values. The PCC is robust to bit depth differences (due to a lack of *L* in Equation S10) and normalisation, as each image is mean-subtracted in its calculation, meaning that it is essentially internally normalised.

If IQMs absolutely must be used for image assessment in microscopy, it is imperative that a) data normalisation is performed and b) the method of data normalisation is fully reported, as it can be seen in Table S2 that the normalisation method can change metric values. The software package and parameters - including those set to default values should also be reported and kept consistent for images being directly compared, as different implementations of metric calculation (especially SSIM) can affect metric values (Table S3).

**Table S2.**
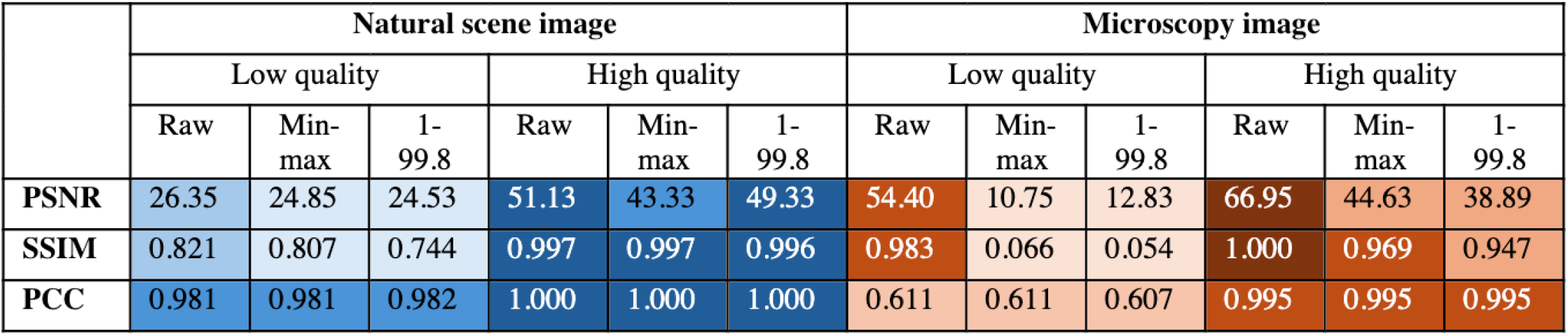
The effect of bit depth and normalisation on IQM values for natural scene and fluorescence images. IQM values are calculated between the reference and low/high quality images displayed in Figure S11. Raw = IQM values calculated without any normalisastion or manipulation of the image bit depth, Min-max = IQM values calculated following min-max normalisation of images (Equation S12), 1-99.8 = IQM values calculated followed 1-99.8 percentile normalisation of images (Equation S11). Table entries are qualitatively shaded according to IQM values (darker = higher values) to highlight the most notable discrepancies.

**Table S3.**
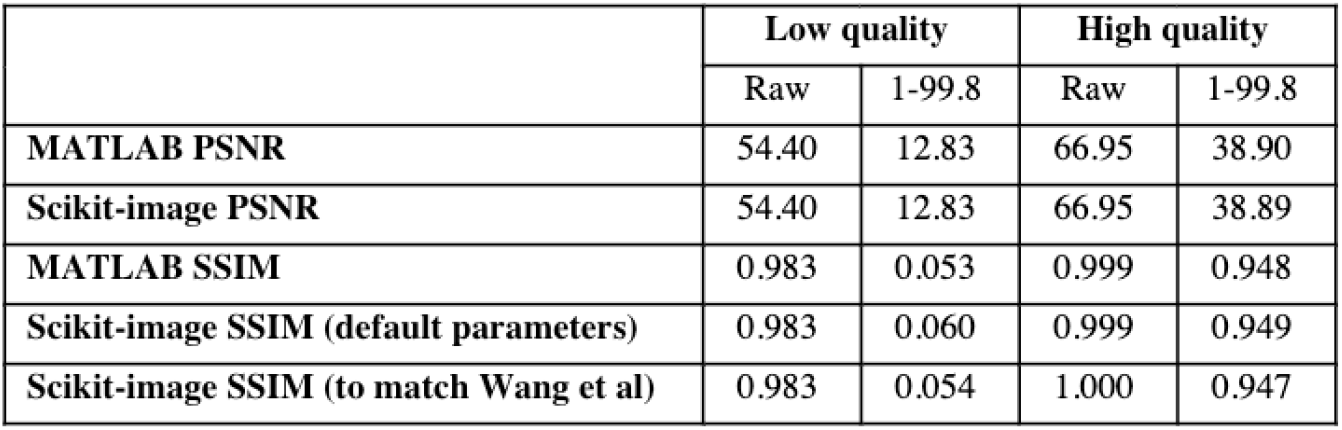
The effect of software implementation on IQMs for fluorescence microscopy data. IQMs were calculated for the low and high quality fluorescence microscopy images against the reference as shown in Figure S11b, with either no data normalisation or 1-99.8 percentile normalisation. MATLAB PSNR and SSIM values were calculated with the `psnr` and `ssim` functions respectively, with no further arguments. Scikit-image PSNR values were calculated with the `metrics.peak_signal_to_noise_ratio` function, explicitly setting `data_range` to 65535 (raw) or 1 (normalised). Scikit-image SSIM (default parameters) was calculated using the `metrics.structural_similarity` function, explicitly setting `data_range` as for PSNR. Scikit-image SSIM (to match Wang et al (1)) was calculated as for defaults, but setting `gaussian_weights=True`, `sigma=1.5` and `use_sample_covariance=False` to match the original implementation of the SSIM method.

**Fig. S11.**
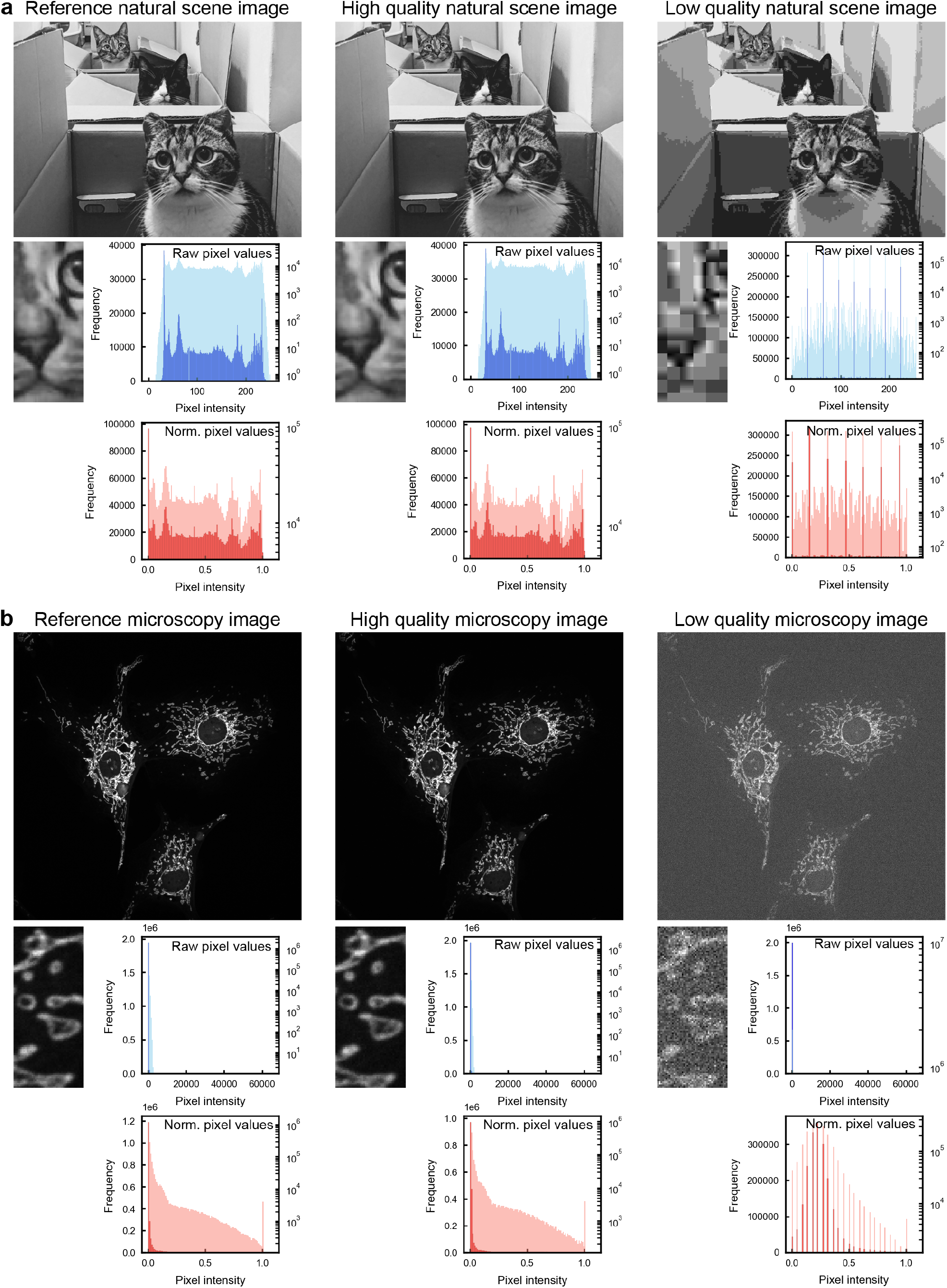
Pixel distributions in natural scene and fluorescence microscopy images. **a** Example of a natural scene images with reference (raw RGB photo converted to 8-bit grayscale), high quality image (JPEG compression level 75) and low quality image (JPEG compression level 1). Inset zooms show loss of detail. Blue histograms show distributions of raw image pixel intensities (dark blue = linear y scale, light blue = log y scale) and red histograms show image pixel intensities following 1-99.8 percentile normalisation (dark red = linear y scale, light red = log y scale). **b** Example of a fluorescence microscopy image (mitochondria in fixed mammalian cells) with reference (illumination dose = 0.21 J/cm^2^), high quality (0.17 J/cm^2^) and low quality (0.007 J/cm^2^) images. Inset zooms show differences in noise. Histograms are displayed as for **a**; here the raw images are 16-bit.

## Notes

### Competing Interest Statement

The authors have declared no competing interest.

