## Supplementary Information for "Image quality metrics fail to accurately represent biological information in fluorescence microscopy"

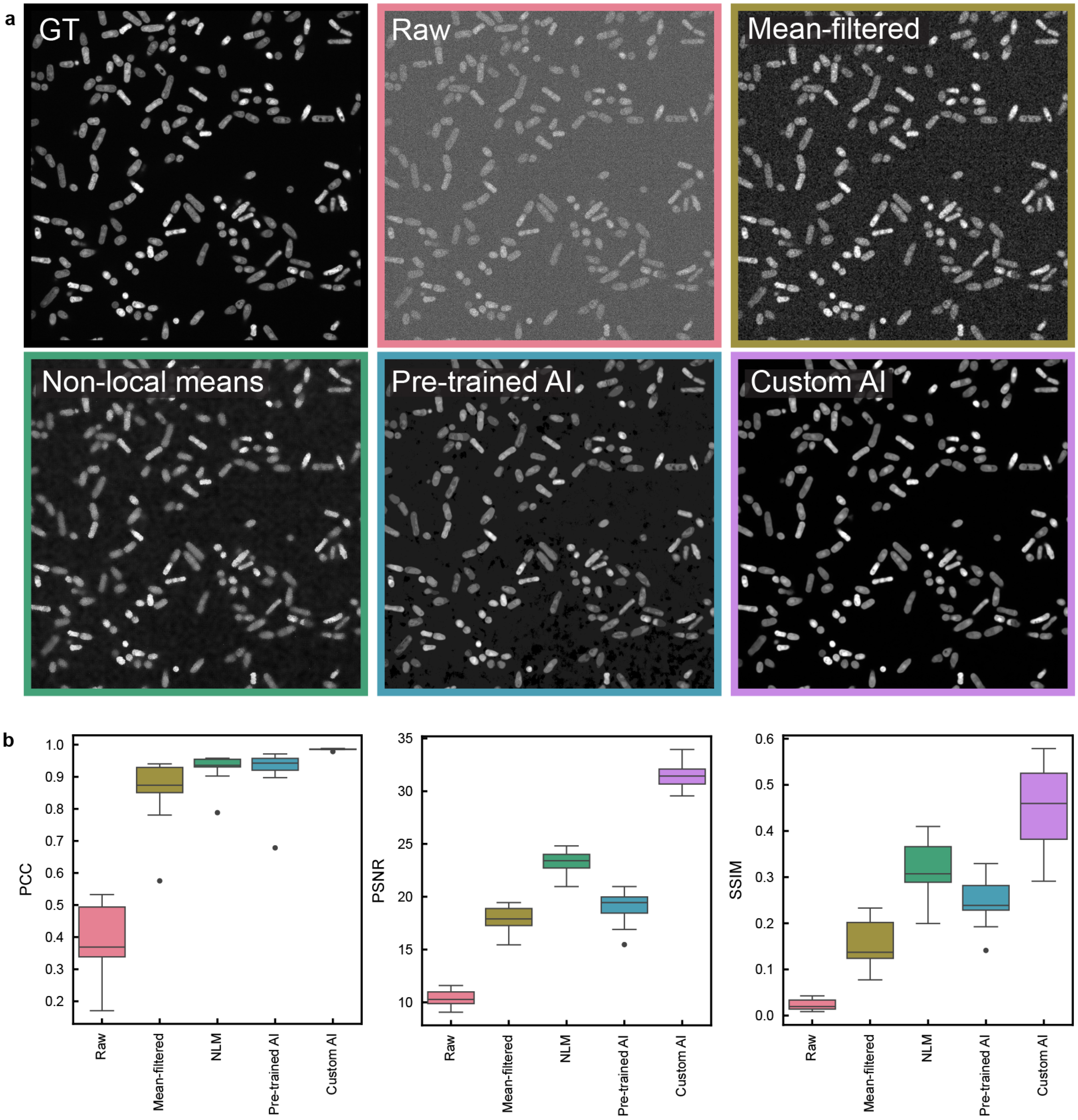


**Fig. S1. Typical application of IQMs to image processing in microscopy. a** Representative images of the same field of view for *S. pombe* expressing sfGFP. Ground truth ('GT') image is acquired at a high illumination dose (0.76 J/cm$^2$), 'Raw' image is acquired at low illumination dose (0.02 J/cm$^2$), and the results of four image processing methods applied to the Raw image are displayed. **b** Box plots showing bulk IQM values for 21 fields of view acquired at 0.02 J/cm$^2$ before (Raw) and after processing. Boxes show three quartile values of distribution, whiskers extend to 1.5 interquartile range. Values outside this range are plotted as individual points.

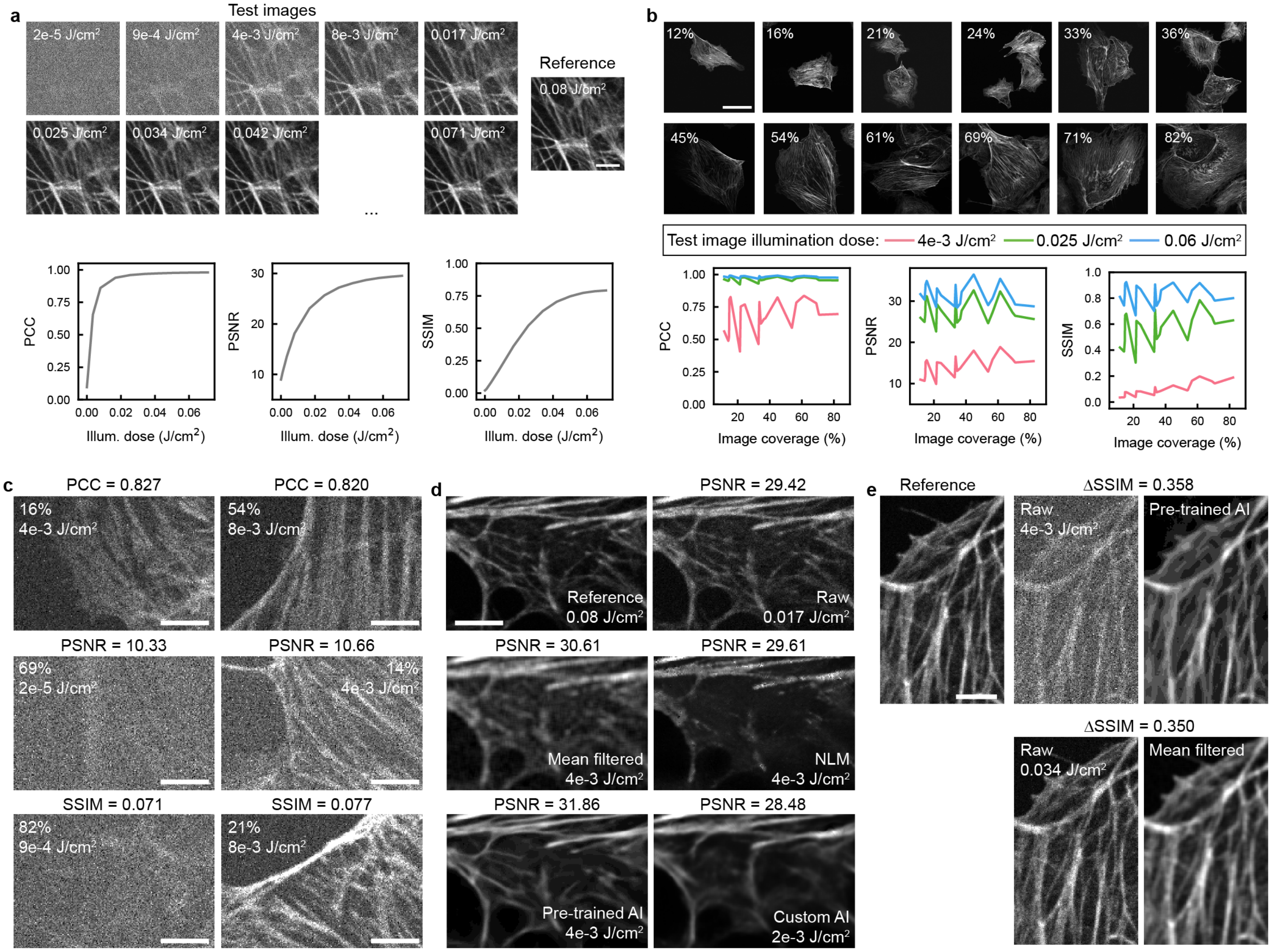


**Fig. S2. IQM behaviour on raw and processed microscopy images of fluorescently-stained actin. a** Top: Crops from a series of images of fixed mammalian cells stained with phalloidin-488. Bottom: IQM values for one whole field of view as a function of illumination dose. **b** Top: Whole fields of view of fluorescently-labelled actin containing different image coverages, acquired at 0.08 J/cm$^2$. Bottom: IQM values plotted as a function of image coverage, for three different test illumination doses. **c** Crops from images where the whole fields of view have a similar IQM value (top: PCC, middle: PSNR, bottom: SSIM) despite acquisition with different illumination doses. **d** Crops from images either raw or following image processing, where each whole field of view has a PSNR $\sim$ 30. **e** Crops from images before and after image processing, where for both images processing has improved the SSIM score by $\sim$ 0.35. In the top panel this corresponds to a substantial improvement in visual quality, whereas in the bottom panel similar features are visible both beofre and after processing. Scale bars: a - 5 µm, b - 50 µm, c - 5 µm, d - 5 µm, e - 5 µm.

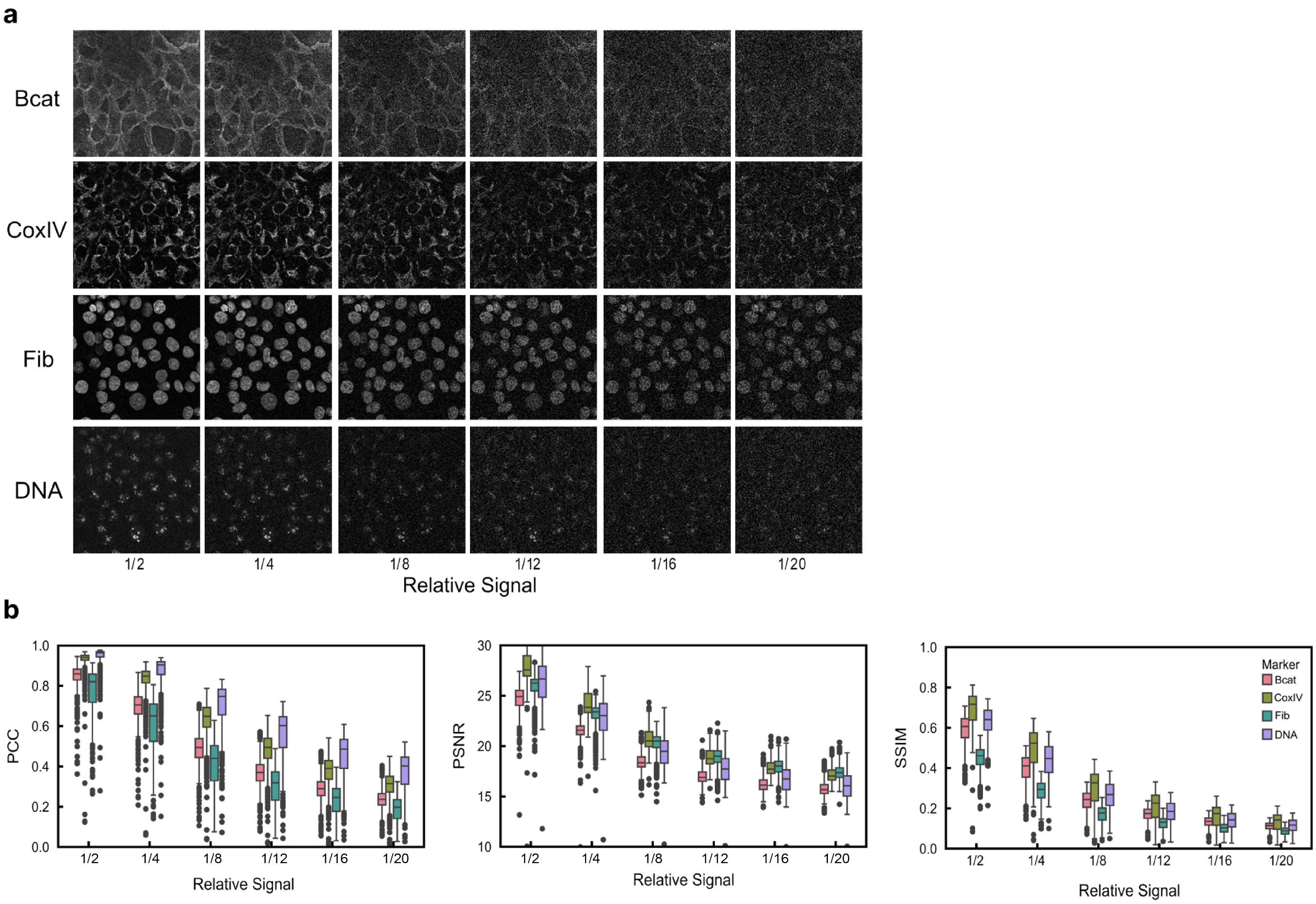


**Fig. S3. Bulk IQM assessment of images for various markers.** **a** Example of image fields for each mark with varying signals (relatively scaled as labelled). **b** Box plots showing range of resulting IQM scores per marker. A clear downward trend observed with less signal. Each box represents scores calculated from 583 different fields. Boxes show three quartile values of distribution, whiskers extend to 1.5 IQR. Values outside this range are plotted as individual points.

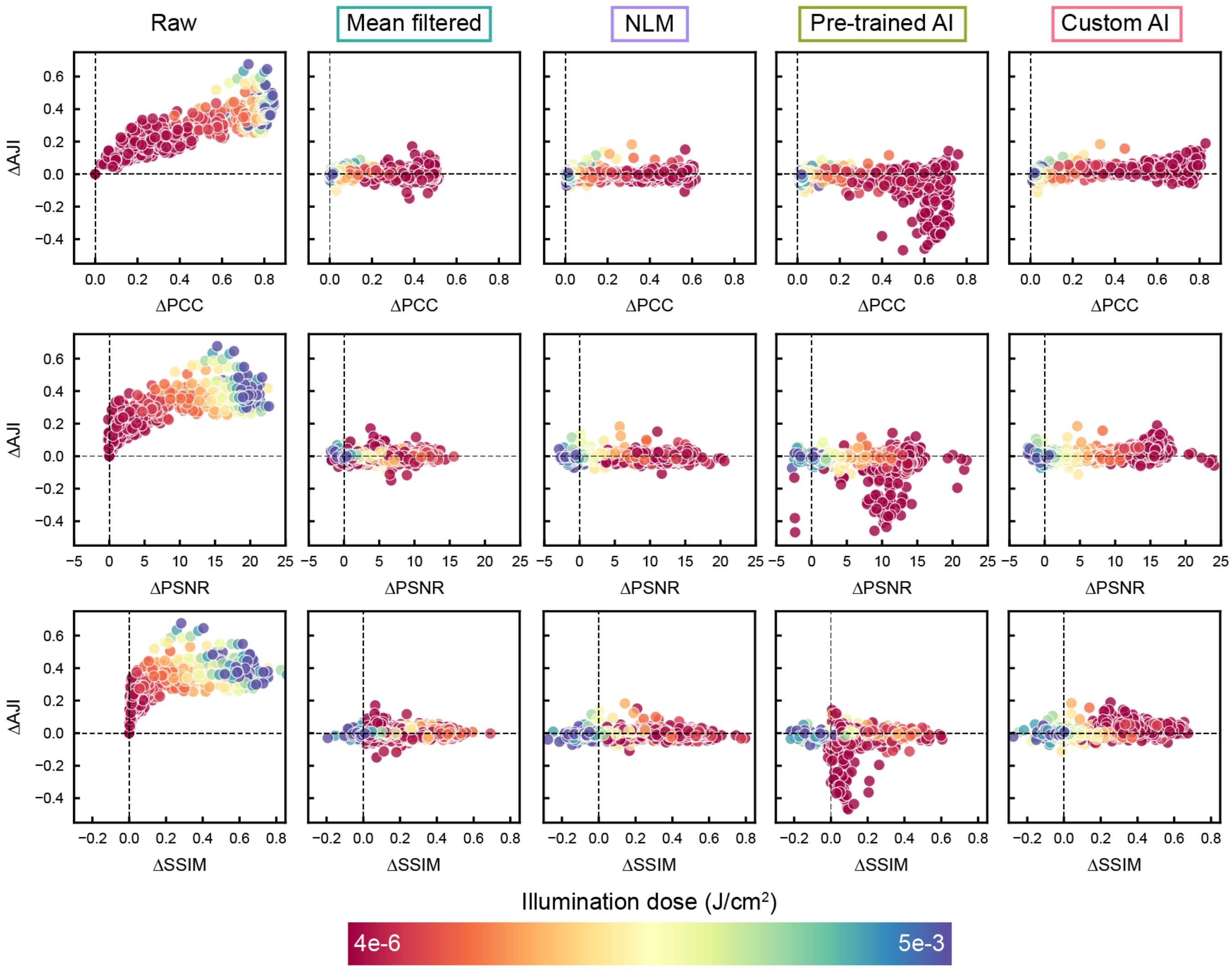


**Fig. S4. Impact of illumination dose on denoising performance and spheroid nucleus segmentation accuracy.** Grouped data from Figure 2c and Figure S7a are split into data points from individual images and colour-coded according to the illumination dose used to acquire the raw image. For the Raw data, ΔAJI and ΔIQM values are calculated relative to the values from the lowest illumination dose for each field of view. For the denoised data, ΔAJI and ΔIQM values are calculated as the difference between the denoised and raw AJI and IQM values for each illumination dose.

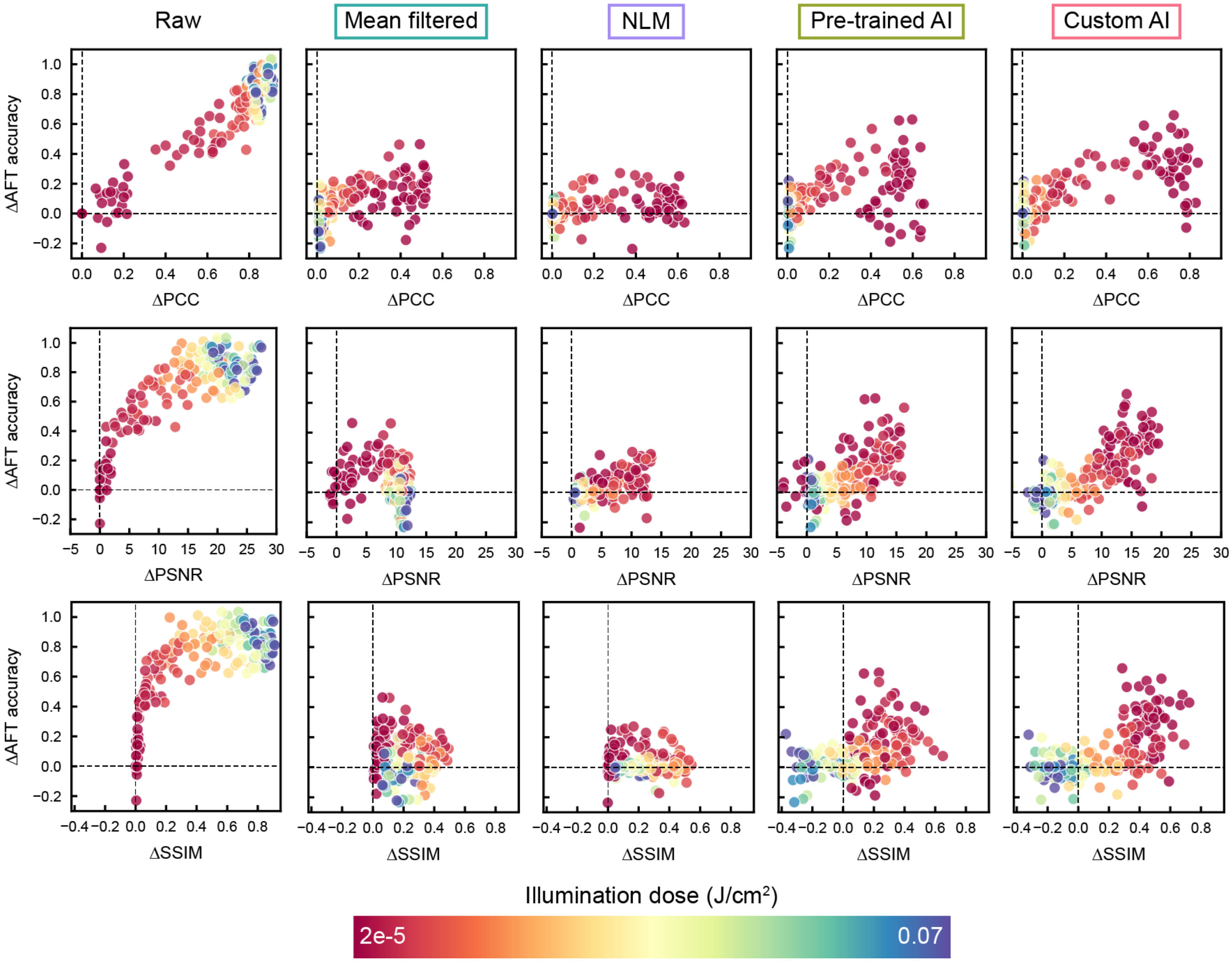


**Fig. S5. Impact of illumination dose on denoising performance and actin filament orientation analysis accuracy.** Grouped data from Figure 2d and Figure S7b are split into data points from individual images and colour-coded according to the illumination dose used to acquire the raw image. For the Raw data, ΔAJI and ΔIQM values are calculated relative to the values from the lowest illumination dose for each field of view. For the denoised data, ΔAJI and ΔIQM values are calculated as the difference between the denoised and raw AJI and IQM values for each illumination dose.

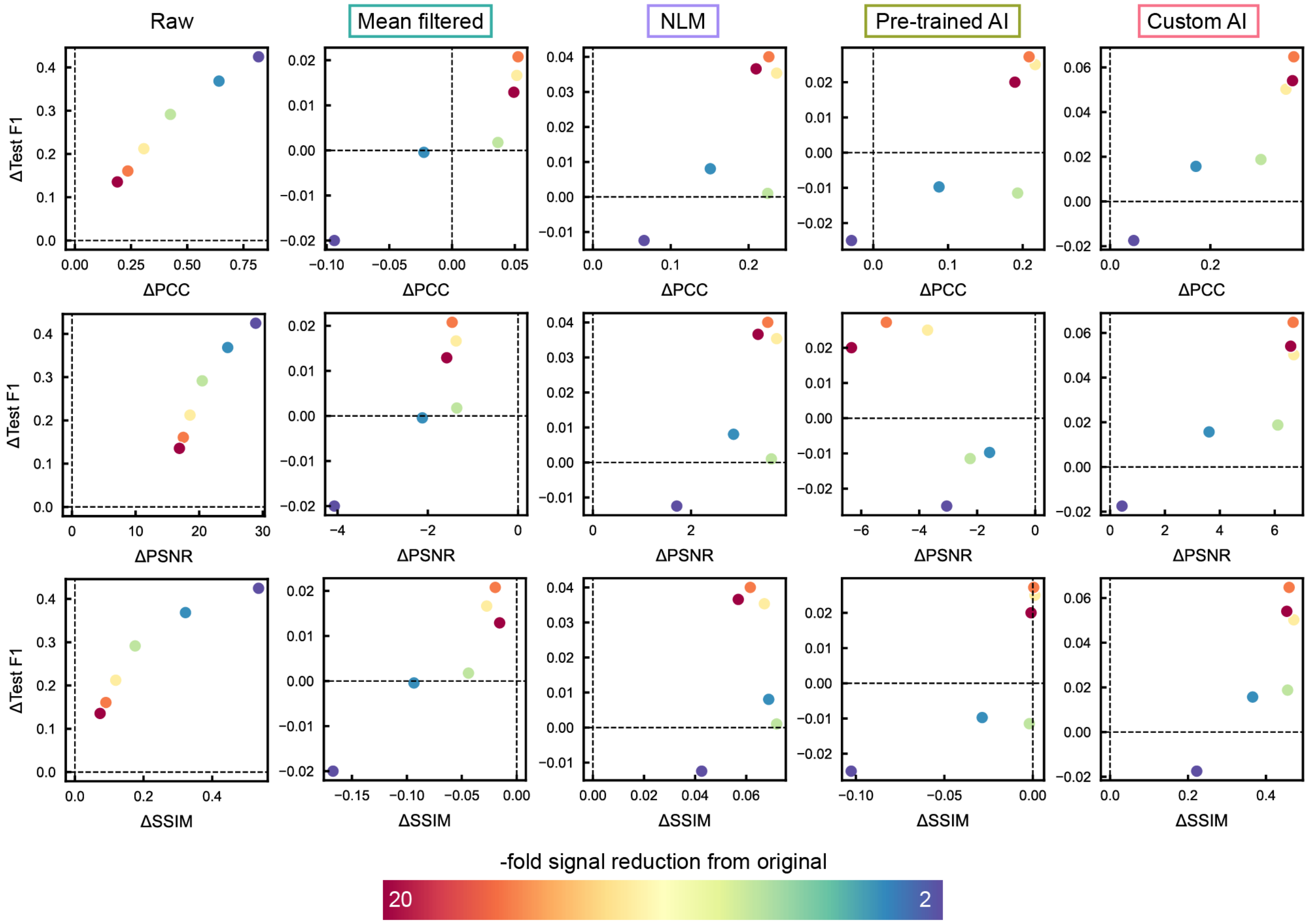


**Fig. S6. Impact of synthetic signal reduction on denoising performance and whole image classification accuracy.** Grouped data from Figure 2e and Figure S7c are colour coded according to the strength of signal reduction applied to generate synthetic noisy images.

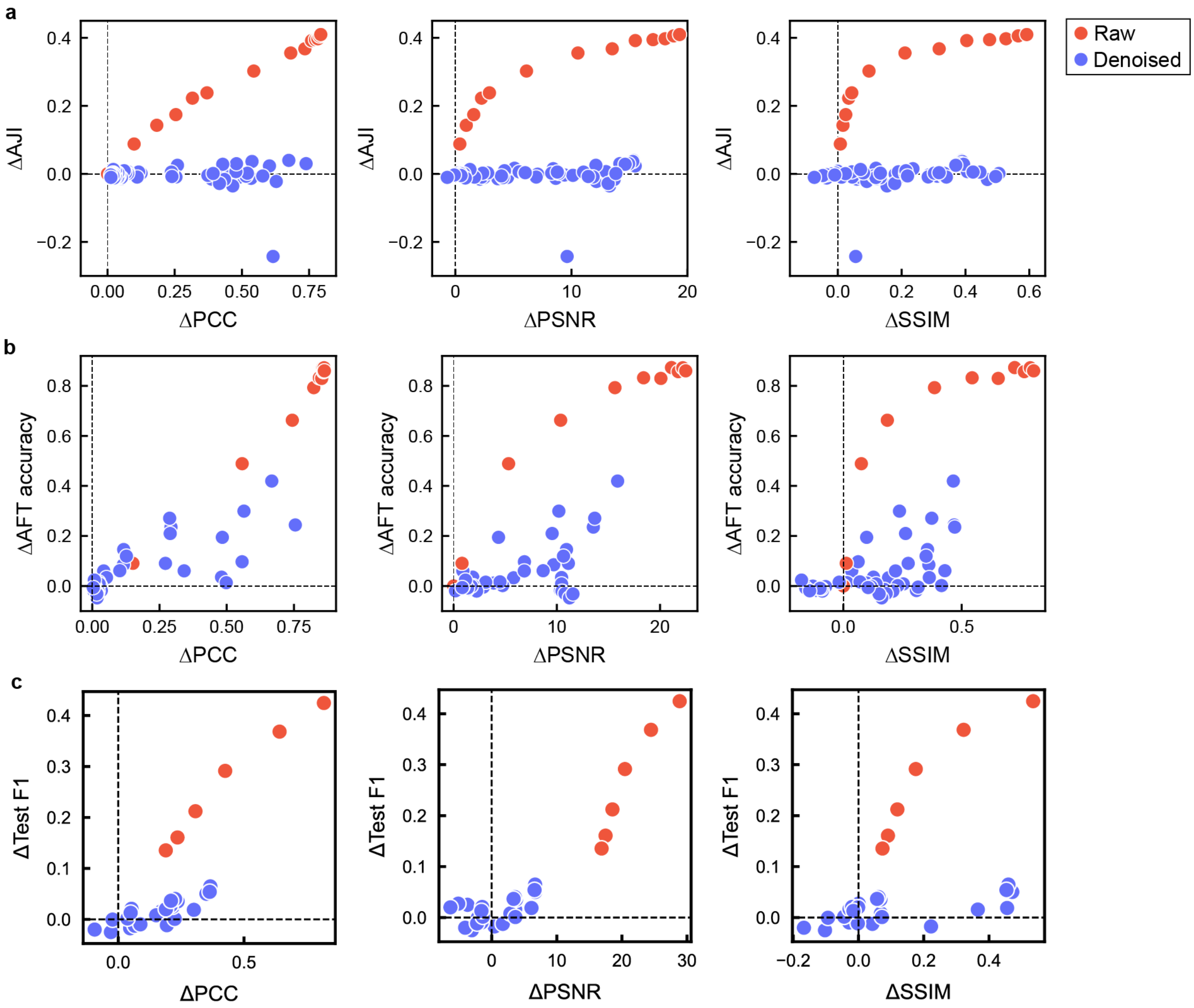


**Fig. S7. Comparison of image analysis accuracy on denoised and raw images. a** Plots of segmentation accuracy (ΔAJI) against ΔIQM for spheroid data. Denoised data are the same as in Figure 2c, but all methods are coloured blue here for clarity. Changes in AJI and IQM value for the raw images are comparisons against the lowest illumination dose image for each illumination dose acquired. **b** As in **a**, but for filament alignment accuracy (ΔAFT). **c** As in **a, b** but for whole image classification accuracy (ΔTest F1). Here, changes in F1 score and IQM value are comparisons against classification performance of reference images (no simulated noise).

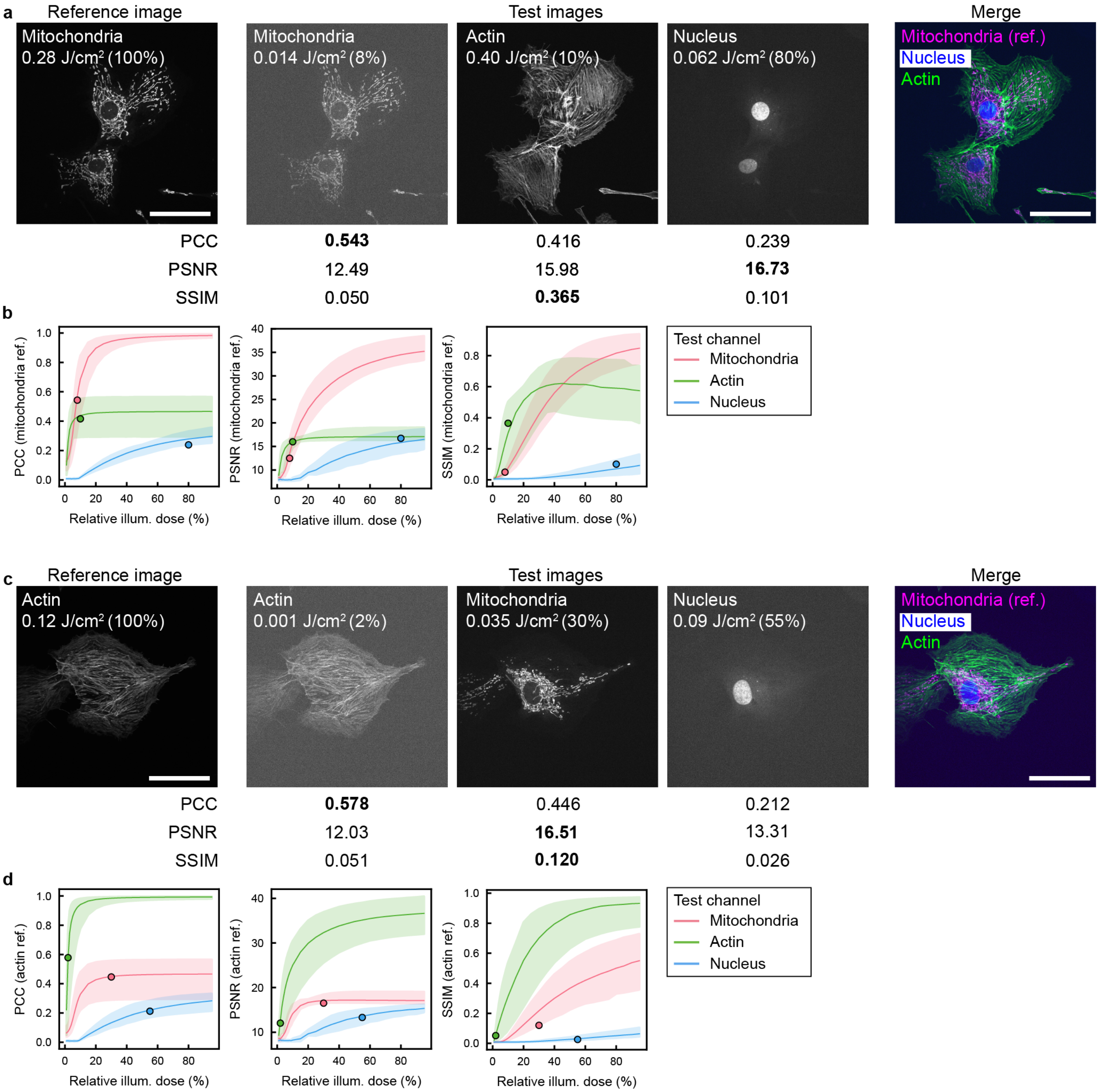


**Fig. S8. Further examples of IQMs 'favouring' incorrect biological structures. a** Images of mitochondria, actin and nucleus stains in the same field of view. The reference image for the IQMs is the mitochondria image at high illumination dose, and the test images are a low illumination dose mitochondria image, medium illumination dose actin image, and high illumination dose nucleus image. Quality metrics are calculated for each image against the reference mitochondria image; numbers in bold indicate the image with the highest scoring IQM. **b** Plots showing the relationship between relative test image illumination dose and IQM for each channel, where in each case the reference is the highest illumination dose mitochondria image for each field of view. Points indicate the images in **a**. **c** As in **a**, but with a high illumination dose actin image as the reference, and low illumination dose actin image, high illumination dose mitochondria image, and high illumination dose nucleus image as the test images. **d** As in **b**, but with the highest illumination dose actin images as the reference, and points corresponding to the field of view in **c**. Scale bars = 50 µm. In all graphs, solid line is mean of N=6 fields of view, and shaded areas extend from minimum to maximum.

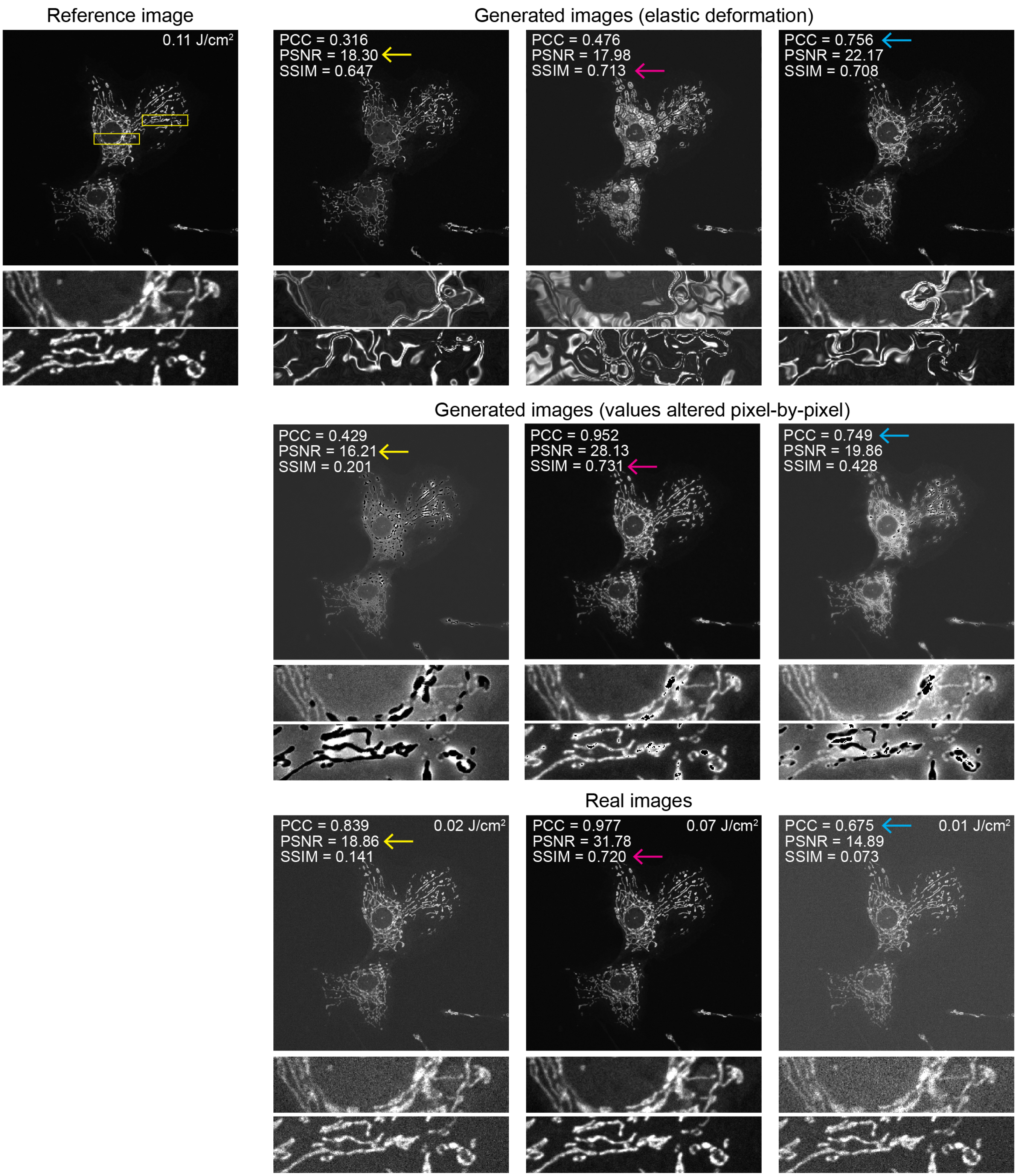


**Fig. S9. Examples of adversarial images obtained via two different image generation strategies and their IQM values compared to real images.** An adversarial attack was performed on various combinations of PCC, SSIM and PSNR metrics (see Figure S10) by optimising an elastic deformation (top row) or pixel-by-pixel value alteration (middle row) of a 0.07 J/cm$^2$ image of mitochondria. IQMs were calculated on 1-99.8% normalised images between the reference image and the generated images. The bottom row shows examples of real acquired images with similar IQM values to the generated images for context (yellow arrows = similar PSNR, magenta arrows = similar SSIM, cyan arrows = similar PCC).

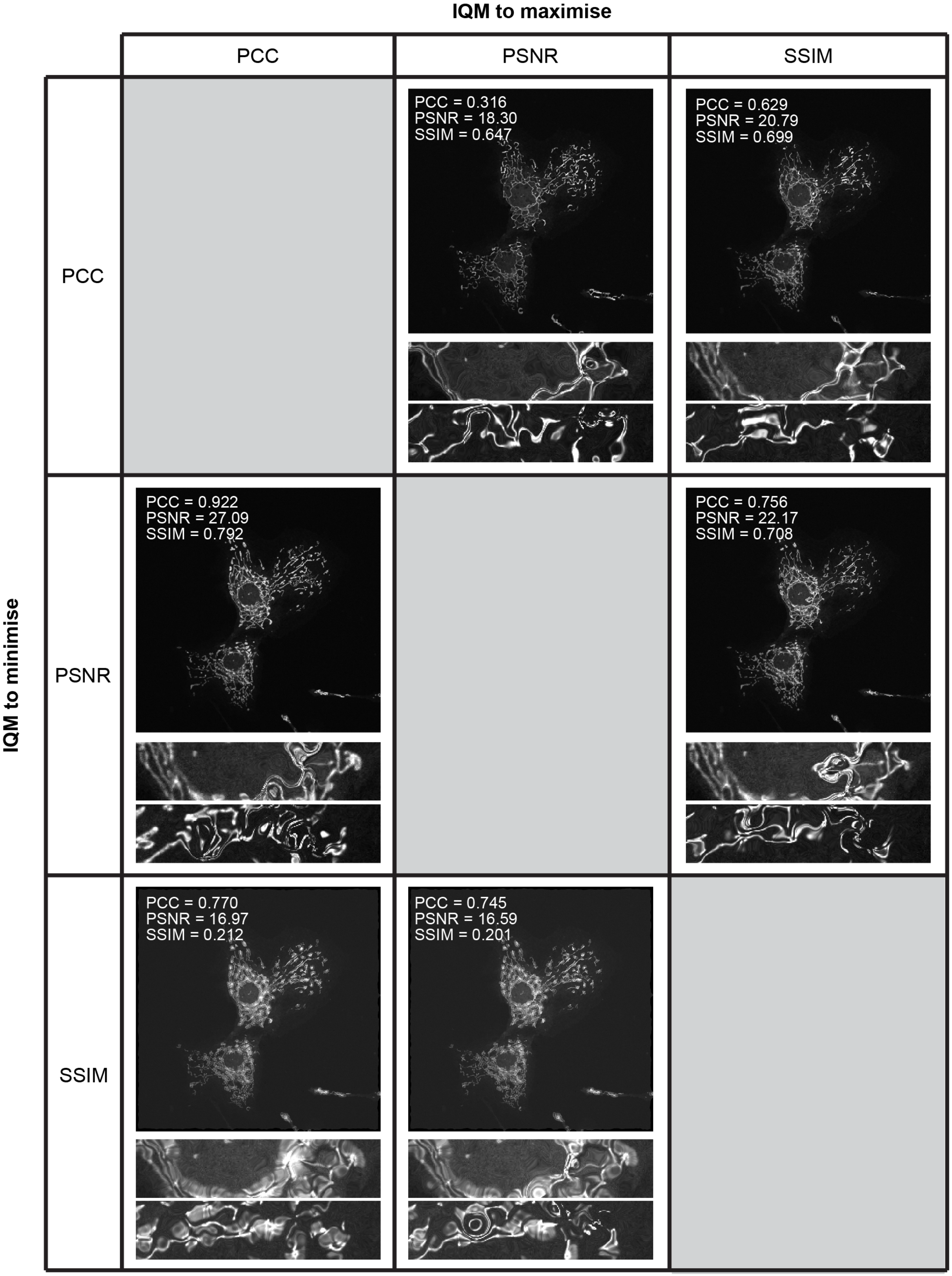


**Fig. S10. Generated images from difference adversarial attack strategies with elastic deformation** Adversarial attacks were performed on the same starting image (mitochondria imaged at 0.07 J/cm$^2$ illumination dose). Columns indicate the IQM which the attack tried to maximise, and rows indicate the IQM which the attack tried to minimise. IQMs were calculated on 1-99.8% normalised images using the 0.11 J/cm$^2$ mitochondria image as the reference (see Figure S9). Zoomed insets are from the same region as shown in Figure S9.

| | | P-values | | |
|---|---|---|---|---|
| Marker 1 | Marker 2 | PCC | PSNR | SSIM |
| DNA | B-cat | 3.17E-123 | 5.78E-59 | 3.66E-18 |
| DNA | CoxIV | 1.08E-06 | 2.22E-36 | 1.36E-39 |
| DNA | Fib | 1.84E-142 | 7.24E-08 | 3.17E-255 |
| B-cat | CoxIV | 2.06E-71 | 3.43E-171 | 1.24E-91 |
| B-cat | Fib | 2.92E-35 | 2.97E-45 | 2.36E-205 |
| CoxIV | Fib | 1.77E-114 | 8.21E-84 | 4.95e-318 |

**Table S1. Significance testing for IQM ranges for different subcellular markers.** P-values are shown for testing the significance of differences between IQM values reported between pairs of markers.

### Supplementary Note 1: Image quality metric formulae, parameters and implementations

**A. Peak signal-to-noise ratio (PSNR).** The peak signal-to-noise ratio is defined as:

$$\text{PSNR} = 10\log_{10}\left(\frac{L^2}{\text{MSE}}\right) \quad \textbf{(S1)}$$

where $L$ is the dynamic range of the image (discussed further below) and MSE is the mean-squared error between the two images, defined as:

$$\text{MSE} = \frac{\sum_{M,N}(X(m,n)-Y(m,n))^2}{M\cdot N} \quad \textbf{(S2)}$$

where $X$ and $Y$ are the reference and test images respectively, each having width $M$ pixels and height $N$ pixels. PSNR originates in signal processing and has been extended to images; there is no reference for PSNR as an image quality metric, however common implementations (for example scikit-image and MATLAB's Computer Vision Toolbox) do not differ in their formulae.

**B. Structural Similarity Index Measure (SSIM).** The structural similarity index measure is defined as (1):

$$\text{SSIM}(X,Y) = \frac{(2\mu_X\mu_Y + C_1)(2\sigma_{XY} + C_2)}{(\mu_X^2+\mu_Y^2+C_1)(\sigma_X^2+\sigma_Y^2+C_2)} \quad \textbf{(S3)}$$

where $X$ and $Y$ are the reference and test images as for the PSNR above. $\mu_X$ and $\mu_Y$ are the mean intensities of the reference and test images, $\sigma_X^2$ and $\sigma_Y^2$ are the variances of the reference and test images, and $\sigma_{XY}$ is the covariance of the reference and test images. The constants $C_1$ and $C_2$ are present to prevent instabilities when $(\mu_X^2+\mu_Y^2)$ or $(\sigma_X^2+\sigma_Y^2)$ are close to zero, and are defined as:

$$C_1 = (K_1L)^2 \quad \textbf{(S4)}$$

$$C_2 = (K_2L)^2 \quad \textbf{(S5)}$$

where $L$ is the dynamic range of the image. Common implementations set the default values of $K_1$ and $K_2$ and 0.01 and 0.03 respectively, as per the original paper. The original paper notes that these values are somewhat arbitrary. The original rationale for developing the SSIM metric was to combine structural, contrast and luminance into a single metric which reproduces key features of the human visual system. Further, the metric was only originally tested on natural scene images and not any biomedical images. Studies in magnetic resonance imaging, for example, have since highlighted discrepancies in SSIM performance and subjective image quality assessment by radiologists (2). Additional parameters are provided in software implementations which are not captured by the SSIM formula. For example, the scikit-image implementation has the option to calculate the SSIM in a window-wise fashion across the image, with further option to spatially normalise the mean and variance of the values within each window by a normalised Gaussian kernel. The MATLAB implementation has the option to change the weighting by which the structure, luminance and contrast terms are weighted within the SSIM formula (presented above with all weights equal to 1). As MATLAB has an option to output an SSIM map, we assume that it also calculates SSIM patchwise across the image, although this is not described in the documentation.

Some microscopy papers use an extension of SSIM, multi-scale (MS)-SSIM (3) as a quality metric. This calculates the luminance, contrast and structure terms of the SSIM metric between the test and reference images with progressive downsampling of the images (typically following Gaussian blurring of the images), and then combines these exponential weights:

$$l(X,Y) = \frac{2\mu_X\mu_Y + C_1}{\mu_X^2+\mu_Y^2+C_1} \quad \textbf{(S6)}$$

$$c(X,Y) = \frac{2\sigma_X\sigma_Y + C_2}{\sigma_X^2+\sigma_Y^2+C_2} \quad \textbf{(S7)}$$

$$s(X,Y) = \frac{\sigma_{XY} + C_3}{\sigma_X\sigma_Y + C_3} \quad \textbf{(S8)}$$

$$\text{MS-SSIM}(X,Y) = [l_m(X,Y)]^{\alpha M}\cdot\prod_{j=1}^{M}[c_j(X,Y)]^{\beta_j}\,[s_j(X,Y]^{\gamma_j} \quad \textbf{(S9)}$$

where $l$, $c$ and $s$ are the luminance, contrast and structure terms respectively. $C_1$ and $C_2$ are defined as above, and $C_3 = C_2/2$. The downsampling scales are represented by $j$, and range from 1 to $M$. $\alpha$, $\beta$ and $\gamma$ are used to adjust the relative contributions of the three terms, and per the original paper parameters are selected such that $\alpha_j = \beta_j = \gamma_j$ for all values of $j$. Furthermore, to normalise cross-scale settings, $\sum_{j=1}^{M}\gamma_j = 1$. The values of parameters were determined in the original paper by asking 8 subjects to compare the quality of 10 sets of natural scene images presented with different degrees of distortion across 5 scales. The obtained parameter values were: $\beta_1 = \gamma_1 = 0.0448$, $\beta_2 = \gamma_2 = 0.2856$, $\beta_3 = \gamma_3 = 0.2363$ and $\alpha = \beta_1 = \gamma_1 = 0.1333$. The authors note that this parameter selection is somewhat crude and that the parameters themselves are "abstract". It should be underlined that the entire rationale of the MS-SSIM metric is, as for SSIM, to mimic the processing power of the human visual system on natural scene images, which is not typically how microscopy images should be assessed.

There is no MS-SSIM implementation provided in the scikit-image python package. Both the pytorch-metrics package and MATLAB have an MS-SSIM implementation where the $k_1$, $k_2$, $\alpha$, $\beta$ and $\gamma$ parameters are set to the values described in the original paper. Both implementations also provide the opportunity to change the number of scales ($M$, with a default of 5), and the sigma of the Gaussian function used to blur the images prior to downsampling (default 1.5 pixels). pytorch-metrics also has an option for normalisation of the MS-SSIM metric, but this is only intended for use when MS-SSIM is being applied as a loss function. When independently demonstrated for use in microscopy, the same parameter values are used as for the original paper and package defaults (4).

**C. Pearson's correlation coefficient.** Pearson's correlation coefficient is calculated as:

$$\mathrm{PCC}(X,Y) = \frac{\sum_{M,N}\left(X(m,n)-\overline{X}\right)\left(Y(m,n)-\overline{Y}\right)}{\sqrt{\sum_{M,N}\left(X(m,n)-\overline{X}\right)^2\sum_{M,N}\left(Y(m,n)-\overline{Y}\right)^2}} \tag{S10}$$

$$I_{\text{norm}} = \text{clip}\left(\frac{I - \text{percentile}(I,1)}{\text{percentile}(I,99.8) - \text{percentile}(I,1) + \epsilon}, 0, 1\right) \tag{S11}$$

where $I_{\text{norm}}$ is the percentile normalised image, $I$ is the original image, percentile$(I,p)$ is the $p$th percentile (pixel value where $p\%$ of the pixels have lower intensity) as calculated using the `percentile` function in numpy, $\epsilon$ is a small constant ($1e-20$) to prevent division by zero, and clip denoised clipping of the pixel range to the range $[0,1]$ to ensure consistency with $L = 1$. For the DU145 and HPA images, which are laser-scenning confocal microscopy data, out-of-distribution pixels should not be produced by the detector and so min-max normalisation was performed as follows:

$$I_{\text{norm}} = \frac{I - \min(I)}{\max(I) - \min(I)} \tag{S12}$$

| | Natural scene image | | | | | | Microscopy image | | | | | |
|---|---|---|---|---|---|---|---|---|---|---|---|---|
| | Low quality | | | High quality | | | Low quality | | | High quality | | |
| | Raw | Min-max | 1-99.8 | Raw | Min-max | 1-99.8 | Raw | Min-max | 1-99.8 | Raw | Min-max | 1-99.8 |
| **PSNR** | 26.35 | 24.85 | 24.53 | 51.13 | 43.33 | 49.33 | 54.40 | 10.75 | 12.83 | 66.95 | 44.63 | 38.89 |
| **SSIM** | 0.821 | 0.807 | 0.744 | 0.997 | 0.997 | 0.996 | 0.983 | 0.066 | 0.054 | 1.000 | 0.969 | 0.947 |
| **PCC** | 0.981 | 0.981 | 0.982 | 1.000 | 1.000 | 1.000 | 0.611 | 0.611 | 0.607 | 0.995 | 0.995 | 0.995 |

**Table S2. The effect of bit depth and normalisation on IQM values for natural scene and fluorescence images.** IQM values are calculated between the reference and low/high quality images displayed in Figure S11. Raw = IQM values calculated without any normalisastion or manipulation of the image bit depth, Min-max = IQM values calculated following min-max normalisation of images (Equation S12), 1-99.8 = IQM values calculated followed 1-99.8 percentile normalisation of images (Equation S11). Table entries are qualitatively shaded according to IQM values (darker = higher values) to highlight the most notable discrepancies.

where $I_{\text{norm}}$ and $I$ are as above, $\min(I)$ is the minimum pixel value in $I$, and $\max(I)$ is the maximum pixel value in $I$. Normalising the microscopy images in Figure S11 prior to calculating IQMs has a significant impact on the PSNR and SSIM values, particularly for the lower quality image (Table S2). In contrast, normalising the 8-bit natural scene images has a less marked effect on IQM values, and percentile normalisation probably has a deleterious effect on metric calculation due to the absence of outlier pixel values. The PCC is robust to bit depth differences (due to a lack of $L$ in Equation S10) and normalisation, as each image is mean-subtracted in its calculation, meaning that it is essentially internally normalised.

If IQMs absolutely must be used for image assessment in microscopy, it is imperative that a) data normalisation is performed and b) the method of data normalisation is fully reported, as it can be seen in Table S2 that the normalisation method can change metric values. The software package and parameters - including those set to default values - should also be reported and kept consistent for images being directly compared, as different implementations of metric calculation (especially SSIM) can affect metric values (Table S3).

| | Low quality | | High quality | |
|---|---|---|---|---|
| | Raw | 1-99.8 | Raw | 1-99.8 |
| **MATLAB PSNR** | 54.40 | 12.83 | 66.95 | 38.90 |
| **Scikit-image PSNR** | 54.40 | 12.83 | 66.95 | 38.89 |
| **MATLAB SSIM** | 0.983 | 0.053 | 0.999 | 0.948 |
| **Scikit-image SSIM (default parameters)** | 0.983 | 0.060 | 0.999 | 0.949 |
| **Scikit-image SSIM (to match Wang et al)** | 0.983 | 0.054 | 1.000 | 0.947 |

**Table S3. The effect of software implementation on IQMs for fluorescence microscopy data**. IQMs were calculated for the low and high quality fluorescence microscopy images against the reference as shown in Figure S11b, with either no data normalisation or 1-99.8 percentile normalisation. MATLAB PSNR and SSIM values were calculated with the `psnr` and `ssim` functions respectively, with no further arguments. Scikit-image PSNR values were calculated with the `metrics.peak_signal_to_noise_ratio` function, explicitly setting `data_range` to 65535 (raw) or 1 (normalised). Scikit-image SSIM (default parameters) was calculated using the `metrics.structural_similarity` function, explicitly setting `data_range` as for PSNR. Scikit-image SSIM (to match Wang et al (1)) was calculated as for defaults, but setting `gaussian_weights=True, sigma=1.5` and `use_sample_covariance=False` to match the original implementation of the SSIM method.

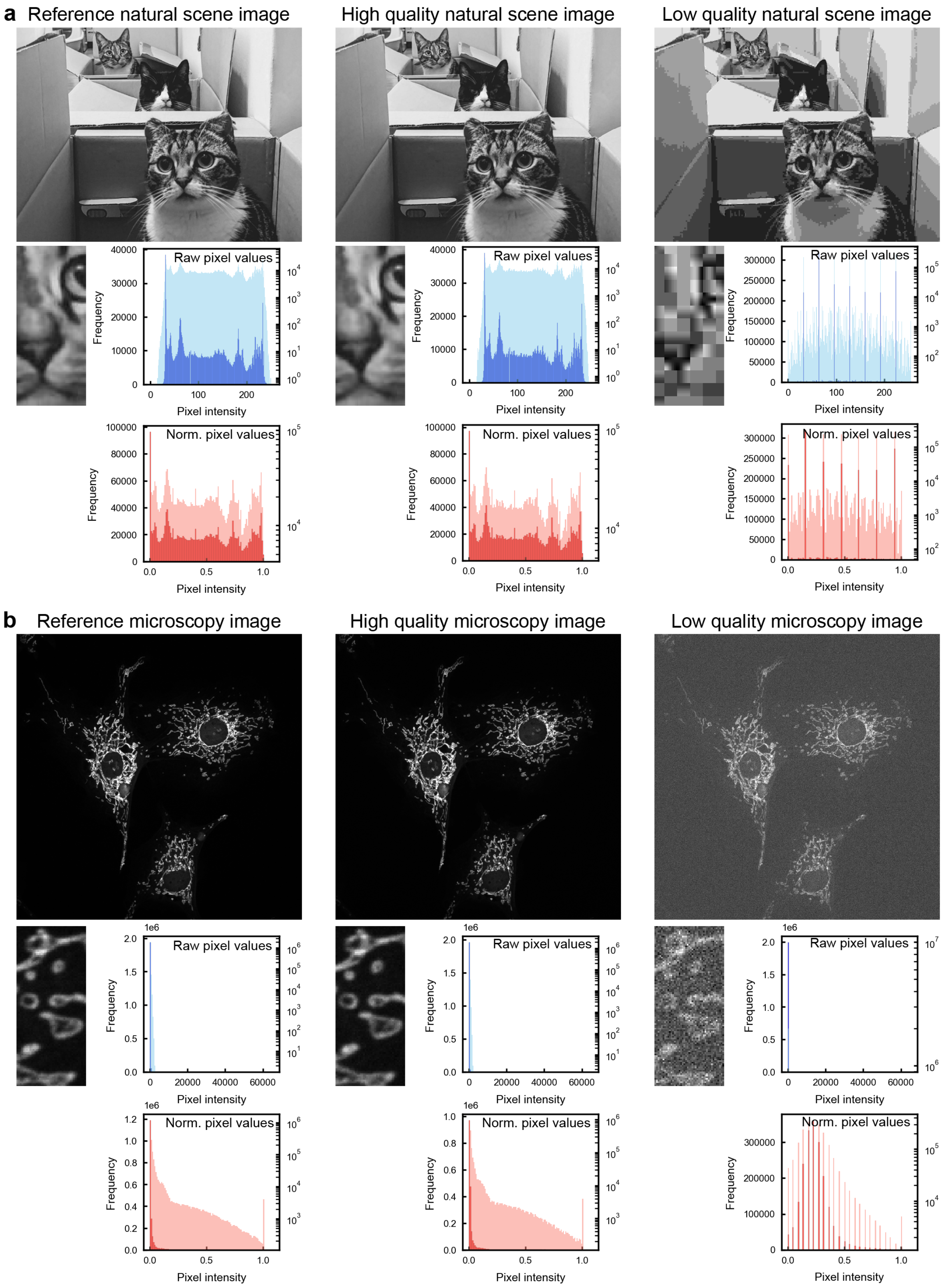


**Fig. S11. Pixel distributions in natural scene and fluorescence microscopy images.** **a** Example of a natural scene images with reference (raw RGB photo converted to 8-bit grayscale), high quality image (JPEG compression level 75) and low quality image (JPEG compression level 1). Inset zooms show loss of detail. Blue histograms show distributions of raw image pixel intensities (dark blue = linear y scale, light blue = log y scale) and red histograms show image pixel intensities following 1-99.8 percentile normalisation (dark red = linear y scale, light red = log y scale). **b** Example of a fluorescence microscopy image (mitochondria in fixed mammalian cells) with reference (illumination dose = 0.21 J/cm$^2$), high quality (0.17 J/cm$^2$) and low quality (0.007 J/cm$^2$) images. Inset zooms show differences in noise. Histograms are displayed as for **a**; here the raw images are 16-bit.
